# Joint Variational Autoencoders for Multimodal Imputation and Embedding

**DOI:** 10.1101/2022.10.15.512388

**Authors:** Noah Cohen Kalafut, Xiang Huang, Daifeng Wang

## Abstract

Single-cell multimodal datasets have measured various characteristics of individual cells, enabling a deep understanding of cellular and molecular mechanisms. However, multimodal data generation remains costly and challenging, and missing modalities happen frequently. Recently, machine learning approaches have been developed for data imputation but typically require fully matched multimodalities to learn common latent embeddings that potentially lack modality specificity. To address these issues, we developed an open-source machine learning model, Joint Variational Autoencoders for multimodal Imputation and Embedding (JAMIE). JAMIE takes single-cell multimodal data that can have partially matched samples across modalities. Variational autoencoders learn the latent embeddings of each modality. Then, embeddings from matched samples across modalities are aggregated to identify joint cross-modal latent embeddings before reconstruction. To perform cross-modal imputation, the latent embeddings of one modality can be used with the decoder of the other modality. For interpretability, Shapley values are used to prioritize input features for cross-modal imputation and known sample labels. We applied JAMIE to both simulation data and emerging single-cell multimodal data including gene expression, chromatin accessibility, and electrophysiology in human and mouse brains. JAMIE significantly outperforms existing state-of-the-art methods in general and prioritized multimodal features for imputation, providing potentially novel mechanistic insights at cellular resolution.

## 1 Introduction

Understanding of molecular mechanisms at cellular resolution provides deeper insights into cellular function, development, and disease progression, but remains elusive. To this end, single-cell multimodal datasets have recently emerged by new sequencing technologies to measure various characteristics of single cells and identify cell functions (e.g., cell types). For instance, patch-seq [1] simultaneously measures gene expression and identifies electrophysiology, and morphological features (beyond omics), e.g., for the mouse visual cortex over 4,000 cells from several brain cell types [2]. [3] profiles single-cell gene expression and chromatin accessibility in the human developing brain. Thus, an integration of single-cell multimodal datasets can significantly aid in our understanding of biological mechanisms contributing to cell types and diseases through the automated discovery of inter-modal relationships.

Many methods have been developed to integrate multimodal datasets to improve prediction of cell types and cellular phenotypes (an outline of the process can be seen in Figure 1a). For instance, our recent DeepManReg [4] approach performs interpretable deep manifold learning which both improves phenotype prediction and is applicable to align single-cell multimodal data. Although these approaches primarily focus on prediction, the underlying idea can be extended to predict data present in separate modalities, referred to as cross-modal imputation going forward (a visual representation can be seen in Figure 1b). Cross-modal imputation is not new, but has been increasingly explored with the advent of deep learning. For instance, BABEL [5] focuses on conversion between RNA and ATAC-seq data matrices through the use of dual autoen-coders, which have the particular advantage of being reusable. Moreover, autoencoders allow for flexibility in latent space formulation. Polarbear [6] utilizes a similar approach, focusing on a conversion network between independently generated latent spaces for each modality. Both methods, however, focus specifically on multi-omics data such as scRNA-seq and scATAC-seq and thus don’t consider high nonlinearity from additional single-cell modalities like electrophysiology. Moreover, they require fully correspondent multimodal data (matched cells), limiting their capabilities with regards to data with missing modalities.

**Figure 1.**
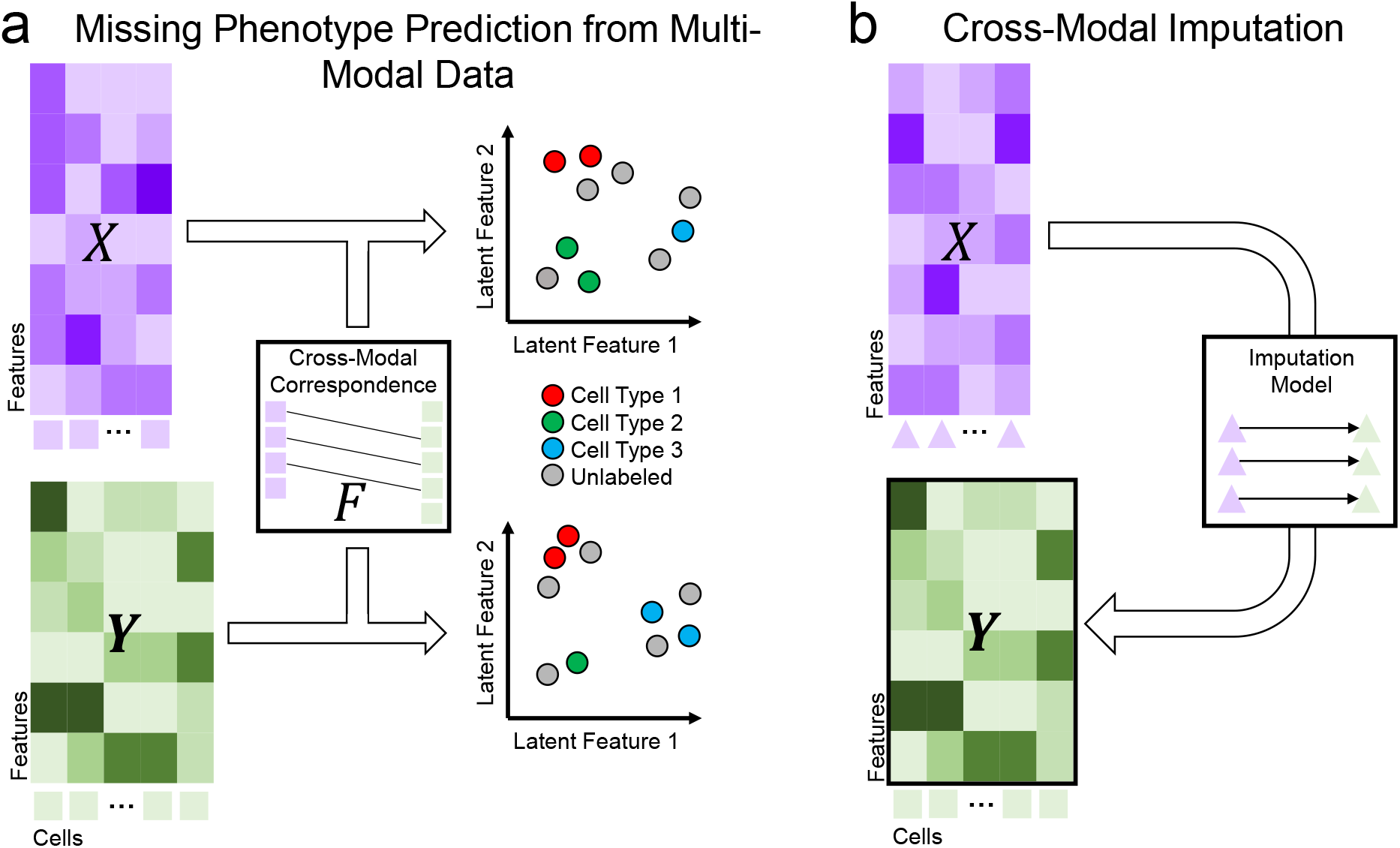
Challenges for multimodal data integration and imputation. (a) Using multimodal data to effectively predict missing phenotypes (e.g. cell types from multimodal single-cell data) is difficult due to heterogeneous features across modalities. Identifying similar latent spaces across modalities enables cross-modal comparison and, by extension, missing phenotype prediction. Machine learning (ML) can be used to discover similar cross-modal latent spaces and enable comparison and phenotype prediction. (b) Certain modalities are cost-prohibitive but may lend significant insight into biological mechanisms. As examples, scATAC-seq data for cell type epigenomics is expensive and electrophysiological data at single-cell resolution is difficult to produce. Imputing one modality from another using ML can alleviate these constraints.

Also, analyzing multimodal datasets incurs additional difficulties, including heterogeneous distributions, multicollinearity, and varying reliability. Several approaches have been thus utilized in aligning multimodal datasets (an outline of the process can be seen in Figure 1b). For instance, Unioncom [7] infers correspondence information from each modality’s distance matrices, then use a modified tSNE method for the final mapping. MMD-MA [8] minimizes an objective function to maximize distribution similarity between datasets while minimizing distortion. ScGLUE [9] uses an autoencoder model to map different modalities onto the same latent space. ScDART [10] learns a latent space and cross-modal relationships simultaneously by predicting gene activity across modalities before projecting to a single latent space. ScAEGAN [11] utilizes dual autoencoders in tandem with cycleGAN [12] to provide aligned latent representations for different modalities. Certain methods rely on user-provided correspondence information to help inform alignment. For example, CLUE [13] assumes completely aligned samples and introduces the impetus for aggregating correspondent latent-space embeddings. ManiNetCluster [14] takes a user-provided correspondence matrix as input and implements nonlinear manifold alignment (NLMA) by solving an eigenvalue problem. ManiNetCluster also implements CCA [15] as a manifold alignment solution for multimodal data. If only partial correspondence information is known, however, existing methods are limiting, and few are designed for such a use-case.

To address these issues, we developed Joint variational Autoencoders for Multimodal Imputation and Embedding (JAMIE). An outline of JAMIE’s capabilities can be found in Supplementary Table 1. JAMIE trains a reusable joint variational autoencoder model to project available multimodalities onto similar latent spaces (but still unique for each modality), allowing for enhanced inference of single-modality patterns [16]. To perform cross-modal imputation, data may be fed into an encoder, then the resultant latent space may be processed by the decoder of the other modality. JAMIE is able to use partial correspondence information. JAMIE combines the reusability and flexible latent space generation of autoencoders with the automated correspondence estimation of alignment methods. We compared JAMIE to state-of-the-art methods on simulation and emerging single-cell multimodal data using gene expression, chromatin accessibility, and electrophysiology in human and mouse brains. We found that JAMIE significantly outperforms other methods (see evaluation in Section 2) and prioritized important multimodal features for multimodal imputation, also providing potentially novel mechanistic insights at cellular resolution.

## 2 Results

As shown in Figure 2a, JAMIE utilizes a joint autoencoder model for data integration and imputation (see details in Section 4. As input, JAMIE takes two data matrices ***X*** and ***Y*** of modalities *X* and *Y* as input. Optionally, an additional correspondence matrix ***F*** may be provided when the samples from two modalities are partially corresponding (e.g. derived from the same single-cells). Encoders in JAMIE transform ***X*** and ***Y*** into latent spaces which are aggregated using available corresponding information. Decoders in JAMIE then predict reconstructions 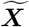 and 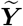 of the original modalities. Please see more details of JAMIE including training, validation, and evaluation in Section 4.

**Figure 2.**
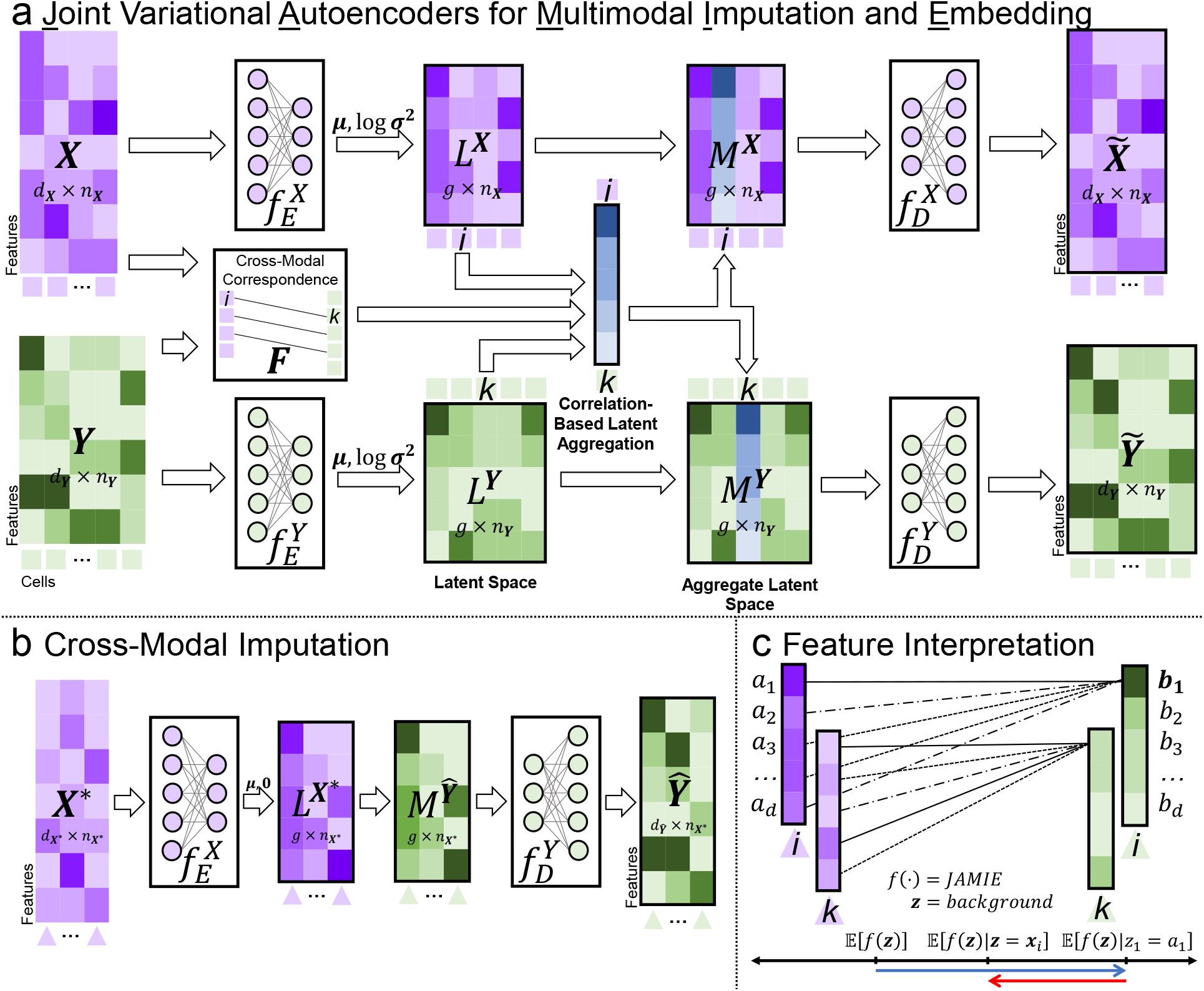
Joint variational Autoencoders for Multimodal Imputation and Embedding (JAMIE) uses variational autoencoders with a novel latent space aggregation technique in order to generate similar latent spaces for each modality. (a) As input, two data matrices ***X, Y*** are provided with optional *user-provided cross-modal correspondence* matrix ***F***. These data matrices are fed through encoding layers 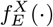, 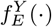 which provide 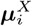, 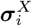 and 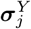, 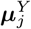 for all samples *i* in ***X*** and *j* in ***Y***, respectively. Then, 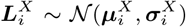 and 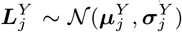 are sampled for all *i* and *j* to produce the latent spaces ***L***^***X***^, ***L***^***Y***^. Latent spaces are aggregated using ***F*** to make ***M***^***X***^, ***M***^***Y***^. Finally, the decoding layers produce a reconstructed version of the original modalities 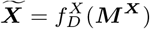, 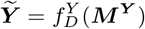 The latent embeddings ***L***^***X***^, ***L***^***Y***^ of the two modalities can be used in tandem for missing phenotype prediction. (b) The trained model can be reused for cross-modal imputation through coupling of encoders and decoders from different modalities. (c) The imputing function 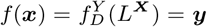, 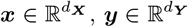 is assessed using Shapley Additive Explanation values [17] which estimate contribution of each input feature by selectively masking the input feature vector with the background.

After training a JAMIE model, its encoder for modality *X* and decoder for modality *Y* can be used sequentially to impute from one modality to another (Figure 2b). Also, the latent spaces from the JAMIE model can be used for phenotype prediction. Furthermore, the use of Shapley Additive Explanation values [17] and similar importance evaluation methods then allows us to prioritize multi-modal features for imputation, as in Figure 2c. These applications are further documented in Subsection 4.12.

### 2.1 Simulated Multimodal Data

We first tested JAMIE on simulated single-cell multimodal data [8]. The simulation data was generated by sampling from a Gaussian distribution on a branching manifold (Figure 3a). We found that the latent embeddings of two modalities in JAMIE preserve the branching structure of the manifold while aligning the cells of the same type in either modality and maintains cell type separation (Figure 3b). To quantify the integration quality, we utilize two metrics: label transfer accuracy (LTA) [7,29], which measures crossmodal phenotype separation, and fraction of samples closer to the true match (FOSCTTM), which measures cross-modal alignment. More details can be found in Subsection 4.9. For separating cell types, JAMIE (LTA= 0.976, FOSCTTM= 0.001) outperforms state of the art alignment methods NLMA (LTA= 0.970, FOSCTTM= 0.001) in LTA and UnionCom (LTA= 0.947, FOSCTTM= 0.079) in both LTA and FOSCTTM (Figure 3c).

**Figure 3.**
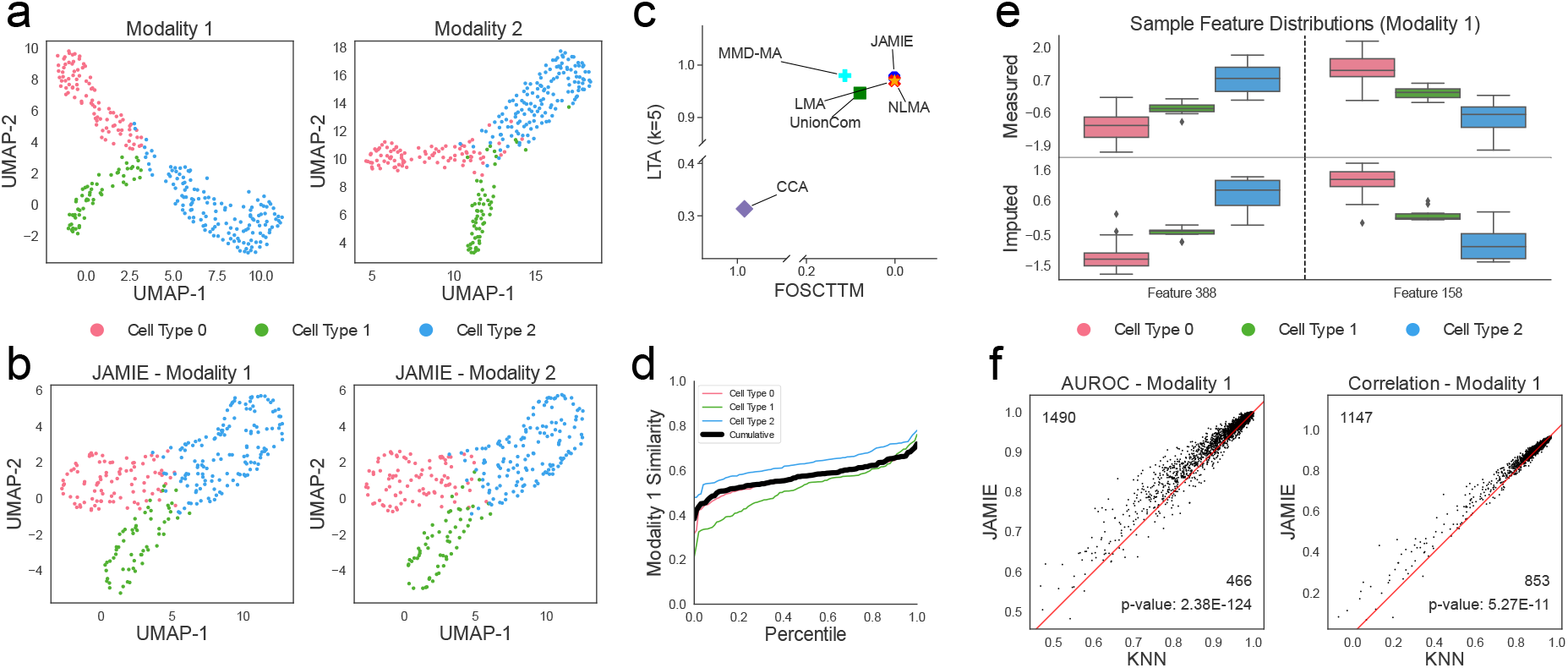
Simulated multimodal data [8]. (a) UMAP of original spaces colored by cell types.(b) UMAP of JAMIE latent spaces. (c) Fraction of samples closer to the true mean (x-axis) and label transfer accuracy (y-axis) of JAMIE and state-of-the-art methods for cell type separation using all correspondence information available. CCA: canonical correlation analysis [15,14], LMA: linear manifold alignment [14], MMD-MA [8], NLMA: nonlinear manifold alignment [14], UnionCom [7]. (d) Cumulative distributions of similarity (1 *−* JS distance) between measured and imputed feature values in Modality 1. The black line indicates the average similarity across cell types while the colored lines each correspond to a single cell type. (e) Measured (top) and imputed (bottom) values of two select features in Modality 1 across cell types (*n* = 300). Boxes span from the upper to lower quartiles and a line indicates the median. Whiskers extend to extremes up to 1.5 times the interquartile range from the upper and lower quartiles. Any outliers beyond this range are plotted individually. (f) Imputation performance for Modality 1 of JAMIE versus a baseline KNN by AUROC and correlation. We utilize a two-tailed binomial test to generate p-values.

Also, we found that the imputed feature values by JAMIE are consistent with the measurements. For instance, as shown in Figure 2d, the imputed values of Modality 1 features have high distribution similarity with the measured values across cell types with average JS distance of 0.428 *±* 0.097 (Figure 3d). Two features with high similarity are highlighted with average JS distances 0.278 and 0.281 (Figure 3e), also showing a preservation of expression changes across cell types (i.e., cell type 1 has a lower value than both other cell types). The imputation performance is compared to a baseline method through correlation and AUROC in subfigure f of each results figure. Each dot represents a cell and the axes are the performance for each method. The red line is the space of equal performance between methods. For imputing the first modality, we also see that JAMIE outperforms the baseline KNN using a two-tailed binomial test for 1, 490 versus 466 features (*p <* 1e-100) in terms of AUROC and 1, 147 versus 853 features (*p <* 6e-11) in terms of correlation (Figure 3f). JAMIE exhibits similar outperformance for AUROC when imputing the second modality (Supplementary Figure 1). We see that JAMIE is able to reliably predict features (See select cells in Supplementary Figure 2).

We tested JAMIE on non-Gaussian simulation data [18] as well and found that JAMIE performed better than the majority of state-of-the-art methods (Supplementary Figure 3). JAMIE achieved an LTA of 0.952 and FOSCTTM of *<* 0.001. The second-best performing method (NLMA) had an LTA of 0.886 and FOS-CTTM of 0.002, similar to the performance of JAMIE with only 75% correspondence information provided (LTA= 0.876, FOSCTTM= 0.016, Supplementary Figure 4). Further details on this dataset can be found in Supplementary Section 2.

### 2.2 Mouse Neuronal Gene Expression and Electrophysiology

We applied JAMIE to infer cross-modal embeddings and impute gene expression and electrophysiological (ephys) data (inhibitory neuronal types: Lamp5, Serpinf1, Sst, Vip, Pvalb, Sncg) neuronal cells in the mouse visual cortex ([2], Figure 4a).

**Figure 4.**
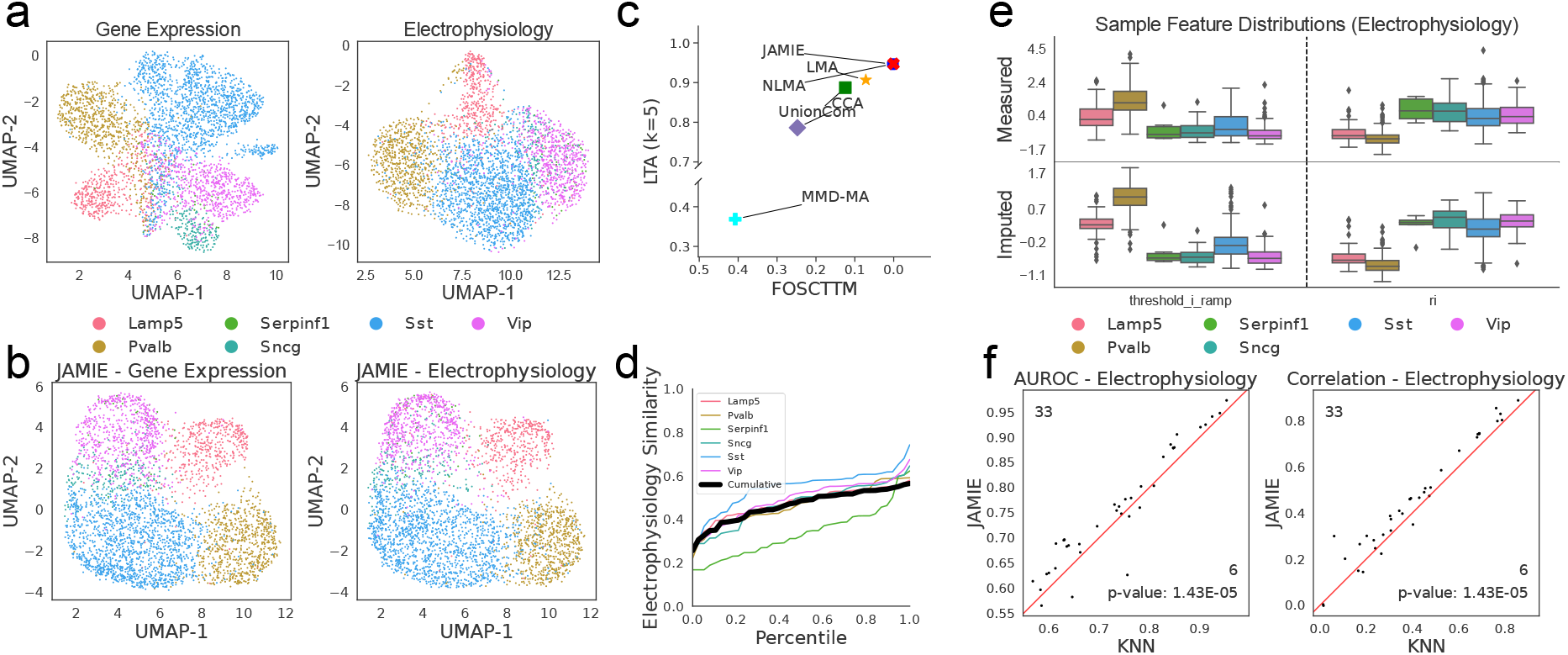
Gene expression and electrophysiological features in the mouse visual cortex [2]. (a) UMAP of single cells by gene expression data (left) and electrophysiological features (right), colored by cell types (inhibitory neuronal types). (b) JAMIE latent spaces of (a). (c) Fraction of samples closer to the true mean (x-axis) and label transfer accuracy (y-axis) of JAMIE and state-of-the-art methods for cell type separation using all correspondence information available. CCA: canonical correlation analysis [15,14], LMA: linear manifold alignment [14], MMD-MA [8], NLMA: nonlinear manifold alignment [14], UnionCom [7]. (d) Cumulative distributions of similarity (1 *−* JS distance) between measured and imputed electrophysiological features by gene expression. The black line indicates the average similarity across cell types while the colored lines each correspond to a single cell type. (e) Measured (top) and imputed (bottom) values of two electrophysiological features across cell types (*n* = 3, 654). Boxes span from the upper to lower quartiles and a line indicates the median. Whiskers extend to extremes up to 1.5 times the interquartile range from the upper and lower quartiles. Any outliers beyond this range are plotted individually. (f) Performance for imputing electrophysiological features of JAMIE versus a baseline KNN by AUROC and correlation. We utilize a two-tailed binomial test to generate p-values.

We found that both gene and ephys embeddings identified by JAMIE can effectively separate those cell types (Figure 4b). Further, JAMIE (LTA= 0.944, FOSCTTM= 0.002) outperforms other alignment methods in both LTA and FOSCTTM, e.g., LMA (LTA= 0.907, FOSCTTM= 0.072) and UnionCom (LTA= 0.887, FOSCTTM= 0.124) (Figure 4c).

Also, JAMIE performs imputation consistently across cell types and generally maintains ephys changes across cell types with an average JS distance of 0.537 *±* 0.115 (Figure 4d). Two ephys features with high similarity demonstrate this preservation with average JS distances 0.314 and 0.316 (Figure 4e). Finally, ephys features are imputed significantly better than the baseline KNN, with 33 of 39 features (*p <* 2e-5) performing better on JAMIE for both AUROC and correlation (Figure 4f). The imputation performance is visualized for select cells in Supplementary Figure 2.

JAMIE was able to generalize to phenotypes beyond cell type, achieving a relatively high LTA (0.650) when predicting cortical layers compared to other state-of-the-art methods (Supplementary Figure 5). For reference, NLMA achieved an LTA of 0.663 while LMA, the next best, only achieved an LTA of 0.515.

In addition to the mouse visual cortex, we also tested JAMIE on gene expression and ephys features in the mouse motor cortex. JAMIE maintains cell type separation in the latent space (Supplementary Figure 6b). JAMIE (LTA= 0.899, FOSCTTM= 0.002) outperforms several methods in both LTA and FOSCTTM, e.g., LMA (LTA= 0.897, FOSCTTM= 0.044) (Supplementary Figure 6c). Unioncom (LTA= 0.248, FOSCTTM= 0.445), CCA (LTA= 0.273, FOSCTTM= 0.366), and MMD-MA (LTA= 0.246, FOSCTTM= 0.478) are unable to align the datasets.

JAMIE also preserves ephys changes across cell types, with average JS distance 0.497 *±* 0.244 (Supplementary Figure 6d). Two ephys features with high similarity demonstrate the preserving nature of JAMIE with JS distances 0.380 and 0.405 (Supplementary Figure 6e). Finally, JAMIE significantly outperforms the baseline KNN using a two-tailed binomial test with 22 of 29 features for correlation (*p <* 9e-3) (Supplementary Figure 6f).

### 2.3 Gene Expression and Chromatin Accessibility in Human Brain

To further investigate the emerging single-cell multiomics data on gene regulation, we apply JAMIE to the gene expression and chromatin accessibility data of the developing human cerebral cortex (scRNA-seq and scATAC-seq by 10x Multiome) [3] (Figure 5a). The chromatin accessibility data measures the accessibility of open chromatin regions (OCRs) by peak signals. OCRs play key epigenomic roles in regulating gene expression.

**Figure 5.**
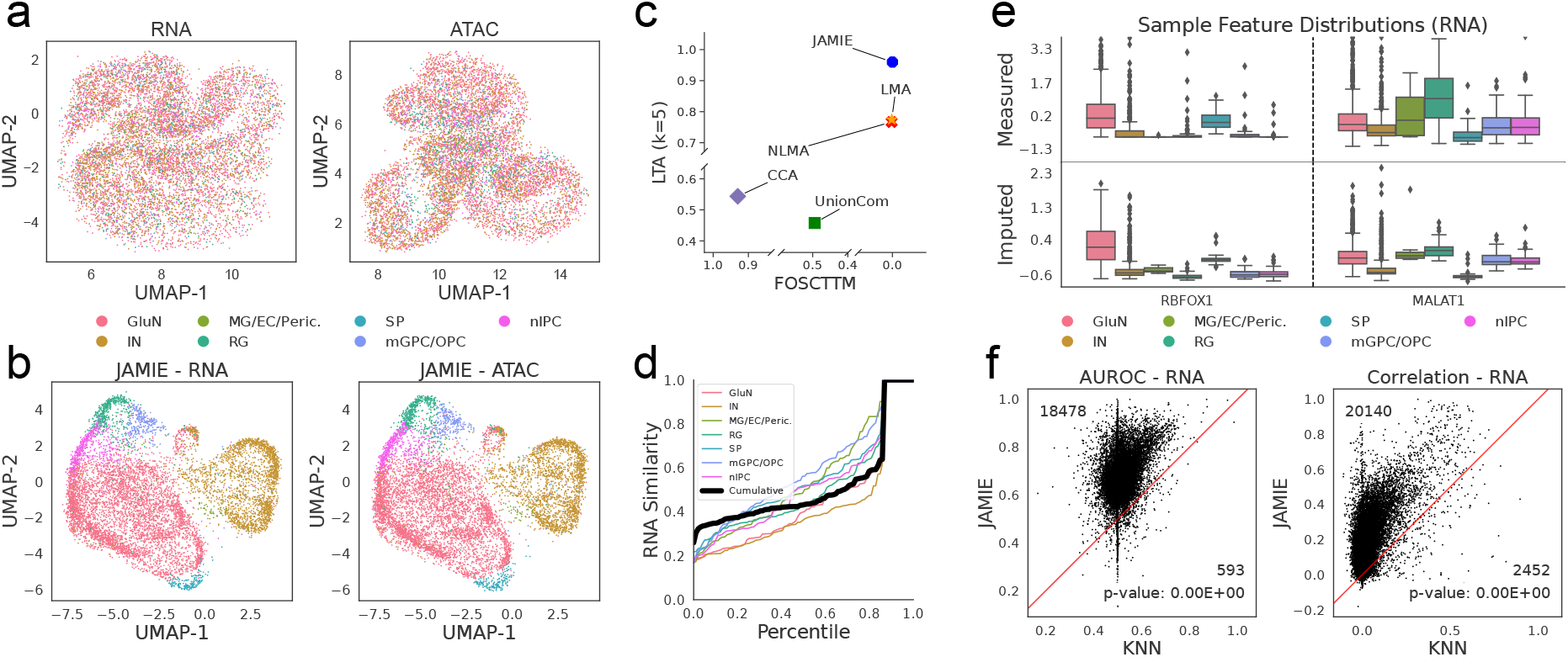
Gene expression and chromatin accessibility of single cells in the developing brain at 21 postconceptional weeks [3]. (a) UMAP of single cells in the human developing brain by gene expression (left) and the accessibility activities of open chromatin regions (right), colored by cell type. (b) JAMIE latent space of a. (c) Fraction of samples closer to the true mean (x-axis) and label transfer accuracy (y-axis) of JAMIE and state-of-the-art methods for cell type separation using all correspondence information available. CCA: canonical correlation analysis [15,14], LMA: linear manifold alignment [14], NLMA: nonlinear manifold alignment [14], UnionCom [7]. (d) Cumulative distributions of similarity (1 *−* JS distance) between measured and imputed gene expression by chromatin accessibility. The black line indicates the average similarity across cell types while the colored lines each correspond to a single cell type. (e) Measured (top) and imputed (bottom) values of two genes across cell types (*n* = 8, 981). Boxes span from the upper to lower quartiles and a line indicates the median. Whiskers extend to extremes up to 1.5 times the interquartile range from the upper and lower quartiles. Any outliers beyond this range are plotted individually. (f) Performance for imputing genes of JAMIE versus a baseline KNN by AUROC and correlation. We utilize a two-tailed binomial test to generate p-values.

JAMIE embeddings separate cell types, in contrast to UMAP alone (Figure 5a-b). In comparison of cell type separations, JAMIE (LTA= 0.959, FOSCTTM*<* 0.001) also outperforms all other methods in both LTA and FOSCTTM compared with NLMA (LTA= 0.767, FOSCTTM= 0.002) and LMA (LTA= 0.775, FOSCTTM= 0.002) (Figure 5c). CCA (LTA= 0.544, FOSCTTM= 0.930) and UnionCom (LTA= 0.458, FOSCTTM= 0.494) failed to align due to the complexity of the data. We note that JAMIE with provided 75% correspondence information (LTA= 0.951, FOSCTTM= 0.047) and 50% correspondence information (LTA= 0.936, FOSCTTM= 0.106) also outperform all other methods in terms of LTA (Supplementary Figure 4).

Moreover, we used JAMIE to impute gene expression from OCRs (peaks) and vice versa. Imputed gene expression values are preserved across cell types with average JS distance 0.484 and standard deviation 0.245 (Figure 5d). We observe wider distributions in the measured data when compared with the imputed data, likely leading to relatively high JS distance with a large number of cell types. We highlight two genes with specifically high similarity, having JS distances 0.260 and 0.332 (Figure 5e). Finally, JAMIE significantly outperforms the imputation baseline using a two-tailed binomial test, better imputing 18, 478 of 19, 071 genes for AUROC (*p <* 1e-100) and 20, 140 of 22, 592 genes for correlation (*p <* 1e-100) from OCRs (Figure 5f). JAMIE exhibits similar imputation performance when imputing ATAC features from RNA (Supplementary Figure 7). Imputation of OCRs from gene expression also outperforms the KNN baseline using the same statistical measure (*p <* 1e-100) (Supplementary Figure 7).

A strong integration method we mentioned in Section 1 is scGLUE [9], which utilizes a combined autoencoder and graph model. To facilitate a comparison between scGLUE and JAMIE, we also ran both methods on [19], which was utilized in [9]. We found that JAMIE (LTA= 0.859 and FOSCTTM*<* 0.001) outperformed scGLUE (LTA= 0.854 and FOSCTTM= 0.038) in both LTA and FOSCTTM (Supplementary Figure 8). NLMA (LTA= 0.259 and FOSCTTM= 0.035) was unable to achieve cell type separation.

### 2.4 Gene Expression and Chromatin Accessibility in Colon Cancer

To further test imputation performance, we compared JAMIE to BABEL [5] using scRNA-seq and scATAC-seq data in colon adenocarcinoma COLO-320DM cells in Supplementary Figure 9 [20]. JAMIE significantly outperforms BABEL in imputing both modalities by a two-tailed binomial test. In particular, JAMIE was better on 12, 309 (*p <* 1e-100) and 13, 334 (*p <* 1e-100) genes by AUROC and correlation, respectively, for gene expression imputation from chromatin accessibility (scATAC-seq, OCR peaks) (Supplementary Figure 9a). For imputing OCRs from gene expression, JAMIE was better on 28, 120 (*p <* 1e-100) and 49, 936 (*p <* 1e-100) features by AUROC and correlation, respectively (Supplementary Figure 9b).

### 2.5 Biological Interpretability for Cross-Modal Imputation

To avoid the black-box nature of many deep learning models, we applied SHAP [17] to prioritize features for cross-modal imputation (Section 4.12). Specifically, this analysis gives top features in one modality for imputing a given feature in another modality.

As shown in Figure 6a, JAMIE prioritized open chromatin regions (OCRs) for imputing the gene, DENND1B, which is a gastric cancer related gene on chromosome 1 [21]. Further, keeping OCRs closer to the location of DENND1B generally results in better imputation performance. For instance, removing OCRs within 10kb produces lower correlation than elsewhere within the chromosome, suggesting the possible capability of JAMIE to reveal the importance of genomic proximity from chromatin accessibility to gene expression.

**Figure 6.**
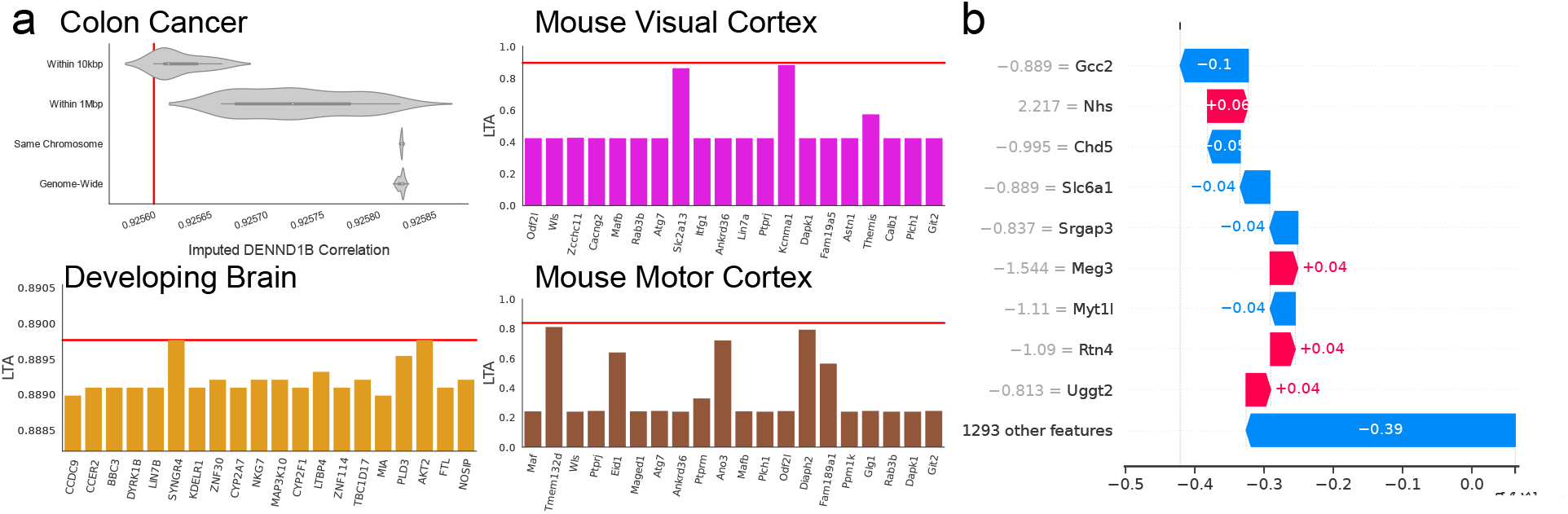
Feature prioritization for cross-modal imputation and embedding. (a) Top left -JAMIE imputation of DENND1B gene with select open chromatin regions removed in colon cancer data [20]. The x-axis is the correlations of the imputed gene expression data with and without select open chromatin regions removed. Aside from the *same chromosome* and *genome-wide* results, all distributions have significantly different means using a one-tailed t-test (All *p <* 2e-10). Top right -LTA for JAMIE with select genes removed in the mouse visual cortex [2]. Bottom left -LTA for JAMIE with select genes removed in the developing brain [3]. Bottom right -LTA for JAMIE with select genes removed in the mouse motor cortex [30]. For each, 14 of the most significant values are displayed along with 6 randomly sampled genes for background. Red Line: Baseline value with no chromatin regions or genes removed. (b) Waterfall plot of select important genes for imputing electrophysiological feature *fast_trough_t_long_square* in mouse visual cortex patch-seq dataset [2]. X-axis: SHAP importance values, Red: Positive SHAP value, Blue: Negative SHAP value.

For the developing brain, JAMIE identifies several important genes, including MIA and BBC3 (Both LTA= 0.889) for contributing JAMIE embeddings to separate the cell types. MIA has been linked to increased risk for neurodevelopmental disorders [22] and BBC3 has been linked to cell death in the adult brain [23]. Additionally, JAMIE identifies the gene SST (LTA= 0.423) as an important gene in the mouse visual cortex. SST is known to be directly related to visual discrimination [24]. Many cell type marker genes were found within the top 200 prioritized genes, with all possible found within the top 400.

## 3 Discussion

JAMIE is a novel deep neural network model for cross-modal estimation. It works for complex, mixed, or partial correspondence multimodal data facilitated by a novel latent embedding aggregation methodology reliant on a joint variational autoencoder structure. In addition to its outperformances as above, JAMIE is also computationally efficient and requires little memory usage (Supplementary Table 2). Moreover, the pretrained model along with learnt cross-modal latent embeddings may be reused for downstream analyses.

We evaluated JAMIE using the evaluation metrics that were designed for the alignment methods such as FOSCTTM and LTA. Despite JAMIE not being explicitly designed to provide common latent spaces in each modality like alignment, JAMIE’s FOSCTTM remains comparable to, and in many applications better than, existing methods in this paper. In addition, LTA appears to be better under complex use cases featuring unfiltered and noisy data. This suggests that the use of modality-specific and aggregated latent spaces in JAMIE allows for flexibility in forming the latent embeddings for better separate cell types from noisy samples.

In cases with low user-provided correspondence, we found that the alignment loss becomes more important. As seen in Supplementary Figure 10a, imputation performance is aided through the inclusion of alignment loss with low (50%) user-provided correspondence. In highly-correspondent applications, the alignment loss may be disregarded to speed up computation by eliminating the computation time of 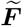 (Supplementary Table 2).

We have demonstrated the capability of JAMIE to impute between two modalities bidirectionally. In practice, imputing a biological outcome from the source modality is likely to be more useful than imputing a source modality from a biological outcome. Examples include imputing electrophysiological features from gene expression and gene expression from epigenomic regulation (e.g., chromatin accessibility). However, our analyses for partial correspondence provide another imputation use case. As stated in our introduction, missing modalities occur frequently when measuring single-cell multimodal data due to its process complexity. This generates data with partial correspondence, i.e., with unmatched cells between modalities. In this case, we showed that JAMIE with partial correspondence is able to outperform a baseline KNN imputation method with full correspondence (using all samples). For instance, as shown in Supplementary Figure 10b, JAMIE imputation using 50% of user-provided correspondence significantly outperforms the baseline by a two-tailed binomial test using all user-provided correspondence while imputing electrophysiological features from gene expression in the mouse visual cortex (*p <* 2e-3).

JAMIE started as a vanilla deterministic autoencoder but was later changed to be variational because of the superior performance of VAEs regarding interpolation in the latent space [25]. Intuitively, the continuous latent space allows for easy sampling and interpolation [26]. Additionally, the proposed aggregation method in Subsection 4.2 relies on the interpolation of latent representations and is improved after switching to a continuous latent space. Though VAEs perform random sampling in the latent space, the process does not add any noticeable randomness to the prediction. Given the same input, the encoder indeed can generate different latent variables due to random sampling during training, but our decoder still reproduces very similar reconstructions. This is because the decoder is trained to map latent samples referring to the same input to very close reconstructions [26]. On the topic of random sampling, the dropout layers do not add randomness to the prediction. They only serve to generalize the model and prevent overfitting by adding redundancy. We chose a default value of *p* = 0.6 to maximize generalizability without adversely affecting performance with default hyperparameters. From Supplementary Figure 11 we see that dropout values of 0.4 and 0.6 have roughly equivalent mean performance while 0.8 has slightly degraded results.

We can split JAMIE into *Preparation* and *Training* phases. The former is preprocessing, PCA, and calculating the inferred correspondence matrix while the latter is the joint VAE model detailed in Figure 2. Supplementary Table 2 shows the consumed time and memory of each phase in seven datasets. For larger datasets, most computation time is spent on the *Preparation* phase. We see that *Training* time scales primarily with cell count while peak memory usage generally scales with both cell count and number of features.

Over the course of runtime, we also found that the KL-divergence dictated the pace of training, as the other losses converge quickly alone (Supplementary Figure 12). Visually, we also see that, as *κ* (Subsection 4.3) increases, an effect can be seen in the alignment and combination losses while the reconstruction loss steadily declines, indicating that the reconstruction and KL losses dominate the training. Without KL-annealing, we found that other losses were unable to converge, even with the KL-loss having a relatively low weight. This was computed on our non-Gaussian simulation data [18].

Training variational autoencoders is time consuming for larger datasets. Thus, prior feature selection such as the automated PCA in JAMIE helps alleviate time requirements. Data preprocessing is also crucial to avoid large or repeated features that can disproportionately shape the features of the low-dimensional embedding, due to the use of reconstruction loss. For cross-modal imputation specifically, the diversity of the training dataset has to be carefully considered to avoid biasing the final model, and negatively affecting the generalizability. JAMIE could also potentially be extended to align datasets from different sources rather than different modalities, such as gene expression measured under different conditions. Because the novel contributions of JAMIE are mainly in the processing of the latent space, many changes in structure and datatype are straightforward to achieve.

## 4 Methods

Figure 2a shows the training process of JAMIE. Two numeric data matrices of modalities *X* and *Y* are used as input. 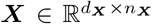 is a data matrix of modality *X* with *d*_***X***_ features and *n*_***X***_ samples. The *i*th column of 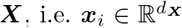, is the feature vector of the *i*th sample of modality *X*. 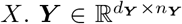 is a data matrix of modality *Y* with *d*_***Y***_ features and *n*_***Y***_ samples. Similarly, the *j*th column of 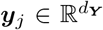, is the feature vector of the *j*th sample of modality *Y*. Each row of ***X*** and ***Y*** is standardized, e.g., *mean* = 0 and *standard deviation* = 1. If available, a user-provided correspondence matrix 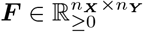 is used to aid construction of the similar latent spaces during training, where ***F***_*ij*_ = 1 implies one-to-one correspondence (i.e. same cells different modalities) between cell *i* in modality *X* and cell *j* in modality *Y*, ***F***_*ij*_ = 0 implies no known correspondences, and 0 *<* ***F***_*ij*_ *<* 1 implies partial confidence of correspondence.

Jamie utilizes joint variational autoencoders to learn similar latent spaces for each of the two modalities, 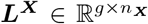 and 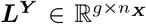, where *g* is the selected dimensionality of the latent space. Higher *g* will lead to better reconstruction loss, but the features will not be as dense information-wise. Lower *g* will condense information more efficiently, but at the cost of reconstruction and imputation performance. We experimentally found *g* = 32 to be robust across multiple datasets, and use this as the default for JAMIE. During training, we utilize correlation-based latent aggregation. In order to learn these latent spaces, JAMIE minimizes the following losses:

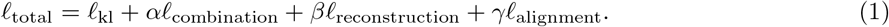

*𝓁*_kl_ takes the Kullback-Leibler (KL) divergence between the inferred distribution of our variational autoencoders (VAEs) and the multivariate standard normal, aiding with the continuity of the latent space. *𝓁*_combination_ enforces similarity of correspondent samples and takes the weight *α. 𝓁*_reconstruction_ is the summed mean squared error between the reconstructions and original data matrices and is assigned the weight *β. 𝓁*_alignment_ uses inferred cross-modal correspondence to shape the resultant latent space and is assigned the weight *γ*. The weights are user-defined and their default values are detailed in Subsection 4.8. An overview of the JAMIE algorithm can be seen in Figure 2.

The default hyperparameters work for a wide variety of cases, and are used for our applications unless otherwise mentioned, but may be customized to the user’s preference.

*𝓁*_kl_ affects the continuity of the latent space and aids in interpolation and stability. This is particularly important for imputation applications. Although this effect is not obvious from the integrated space itself (Supplementary Figure 13a), the imputation performance is significantly improved with this loss (Supplementary Figure 13b).

*𝓁*_combination_ controls the focus on similarity between correspondent features across modalities. A higher weight will result in closer cross-modal cell representations at the risk of losing per-modality detail. *𝓁*_reconstruction_ controls the ability of the latent space to encode raw information from each modality. A larger weight will make cell representations more adherent to their single-modal representations at the cost of cross-modal representation similarity of correspondent samples. The effects of these losses are very pronounced when removed from the training process, and result in lower integrated space reliability (Supplementary Figure 13a).

Finally, *𝓁*_alignment_ is primarily helpful for low-correspondence applications where the shape of the latent space is not clear from context. A higher weight will increase the similarity of cross-modal correspondent cell representations by changing the shape of the resultant latent space. This can come at the cost of cell type mixing and cluster accuracy but can result in better imputation performance in low-correspondence applications (Supplementary Figure 10a).

The latent dimension *g* can be lowered for more compressed representations, potentially at a loss in accuracy (Supplementary Figure 14a). This can be useful in applications which present overfitting, normally when the dataset has very few features or samples. The number of principal components used in the JAMIE model can be adjusted to change model size, which can speed up calculation at the cost of performance (Supplementary Figure 14b). This may also be lowered to avoid overfitting.

### 4.1 Distribution of Latent Space Features

Intermediate feed-forward neural networks are utilized to generate distributions for each feature in the *g*-dimensional latent space for each modality. These will be henceforth referred to as encoders, denoted 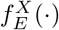, 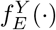 for each modality. The variational encoders have four hidden layers. The encoder of modality *X* transforms the feature dimension as follows: 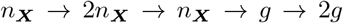. For modality *X*, the top half of the fourth layer output corresponds to means and the bottom half corresponds to log variance per sample and feature as in Equation 2a. The encoder is similar for *Y*. Batch normalization and leaky ReLU activations are included wherever possible, and dropout layers are added (*p* = 0.6) for imputation models.

For each sample ***x***_*i*_ and ***y***_*j*_, we sample the multivariate Gaussian

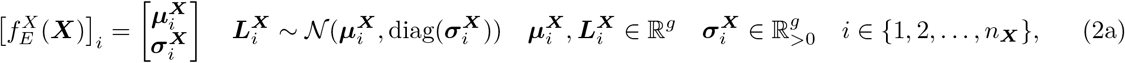

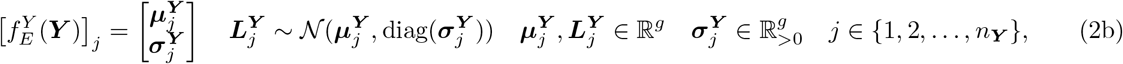

where 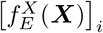 and 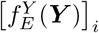 are the encoded results of ***x***_*i*_ and ***y***_*j*_, respectively, and diag(·) is a diagonal matrix generated from a vector. 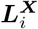 and 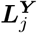 are sampled from the multivariate Gaussian distributions indicated above. We define 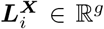 as the *i*th column of ***L***^***X***^, representing the feature vector of the *i*th sample. 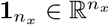 is defined similarly.

The VAE is able to approximately map any non-Gaussian input data onto the multivariate Gaussian latent space (using the encoder) and reconstruct the input data from the space using the decoder [26]. Therefore, the multimodal input to JAMIE is not restricted to a Gaussian distribution.

### 4.2 Aggregation of the Latent Spaces

Using our user-provided correspondence matrix 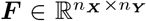, we can perform an aggregate calculation to combine the latent space vectors of known aligned points. In particular,

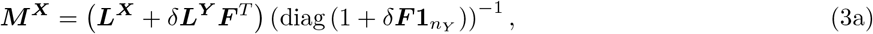

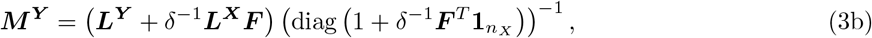

where the aggregate feature vector 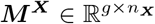 has *g* features and *n*_***X***_ samples and 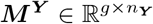 has *g* features and *n*_***Y***_ samples. 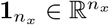 and 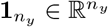 are vectors of all ones. *δ* refers to the relative weighting between modalities *X* and *Y* in the aggregation function. We learn *δ* over the course of training. A value *δ <* 1 implies that ***X*** is weighted more than ***Y*** during aggregation, while *δ >* 1 implies the opposite. The construction of ***M***^***X***^, ***M***^***Y***^ is shown visually in Figure 2a in the formulation of the blue vector. Figure 2a shows the case ***F***_*ik*_ = 1 with all other entries in row *i* and column *k* being 0, which results in simply averaging the latent feature vectors of ***x***_*i*_ and ***y***_*k*_.

The linear aggregation technique is backed by the nonlinear transformation of the encoder, which is able to account for varying timings, magnitude changes, and distributions due to its deep structure. In Figure 2, this is the blue latent space vector combination of correspondent samples. We term the technique *correlation-based latent aggregation*. This moves correspondent latent embeddings in similar directions over the course of training and is key in the formation of similar latent spaces. Correlation-based latent aggregation is the primary motivation for the use of the variational autoencoder framework, as it allows for this intuitive representation of partially aligned datasets. We can now adjust our low-dimensional data embeddings by using aggregates wherever possible to produce our final aggregate latent spaces ***M***^***X***^ and ***M***^***Y***^.

### 4.3 Continuity of the Aggregate Latent Spaces

Continuous ***L***^***X***^ and ***L***^***Y***^ facilitate imputation, thus, we adopt the VAE architecture [25] and the corresponding KL loss:

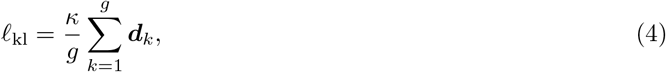

where ***d***_*k*_ takes the KL divergence of the *k*th feature between the distribution of our latent space estimate and a multivariate standard normal as the targeted distribution:

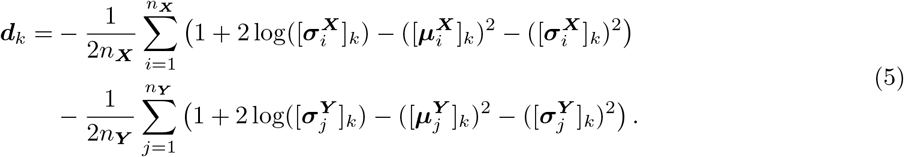

The parameter *κ* is slowly increased from 0 to 1 over the course of training to accelerate convergence [27], further detailed in Subsection 4.7. This process is further referred to as *KL-annealing*. KL divergence is described in Subsection 4.7.

*𝓁*_kl_ will lower to zero as the latent space’s distribution approaches the standard multivariate Gaussian. This encourages a continuous latent space through the nonzero standard deviations. The mean approaching zero encourages a small, clustered latent space. This will negatively affect metrics such as silhouette distance, but can aid in imputing outlier or unseen single-cell data.

### 4.4 Similarity of the Original and Aggregate Latent Spaces

The structure alone with JAMIE already produces shared latent spaces ***M***^***X***^ and ***M***^***Y***^. To balance ***L***^***X***^ and ***L***^***Y***^ and keep both latent spaces the same scale, we implement a loss proportional to the square norm of the difference between each latent space and its aggregated matrix ***M*** :

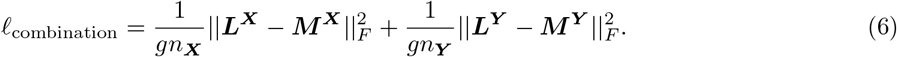

The combination loss makes the aggregate matrices as similar as possible to the originals while still maintaining cross-modal corresponding cell representation similarity. In practice, as long as the two modalities contain unique information and are non-trivially corresponding, the combination and reconstruction losses will balance each-other out, as the reconstruction loss ensures that information unique to each modality is encoded in the latent representations. This balance allows JAMIE to form the latent space flexibly while preserving as much single-modal detail as possible.

### 4.5 Similarity of the Original and Reconstructed Data Matrices

Decoders are structured in the reverse manner as encoders. It uses fully connected layers to transform the feature dimension as follows: 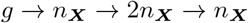. The structure is similar for *Y*. Batch normalization and leaky ReLU are included where possible, and dropout layers are added (*p* = 0.6) for imputation models. We denote the decoders 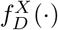, 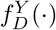 for each modality and use each to get an estimate of the original datasets 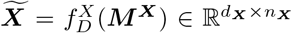 and 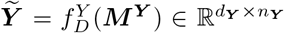. The quality of the reconstruction, and thereby the information retained in the latent space, can be evaluated by:

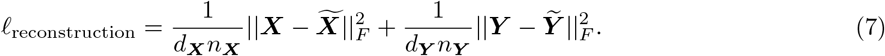

### 4.6 Correspondence of the Aggregate Latent Spaces

We find the optimal inferred correspondence matrix 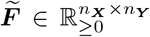by solving the following optimization problem:

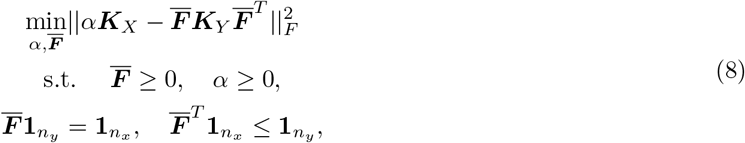

where [***K***_*X*_, ***K***_*Y*_] = [*dist*(***X***), *dist*(***Y***)] are the intra-dataset distance. This generally uses geodesic distance and is similar to the quadratic assignment problem.

The procedure originates from an extension on [28], which optimized a similar expression but assumed one-to-one correspondence on inter-dataset cell pairs. [7] extended this to the concept of one-to-many correspondence between modalities and created the implementation in Equation 8.

Once solved, 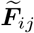 denotes the probability that sample ***x***_*i*_ is matched with sample ***y***_*j*_. The objective of 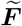 is to provide complete correspondence information. ***F*** will generally contain only one-to-one relationships, while true inter-modal relationships are often layered and complex. In practice, we may consider combining ***F*** and 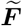 by a weighted average if the user-provided correspondence is especially low (see Algorithm 1).

To ensure that correspondent samples have similar aggregate latent representations we introduce *𝓁*_alignment_. This metric relies on our inferred correspondence 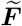 rather than the user-provided correspondence matrix ***F***. Unlike ***F***, our inferred correspondence 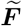 contains information about samples which are likely to not correspond. We can utilize this dense information to shape our latent space such that contributing samples are clustered:

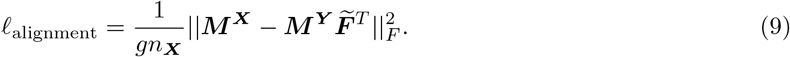

The alignment loss is primarily effective when the user-provided correspondence is weak (containing few highly-correspondent pairs). This loss shapes the integrated latent spaces and aids in obtaining cross-modal latent similarity by inferring correspondence using unsupervised manifold alignment. Any unsupervised correspondence method may be substituted for 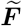.

### 4.7 Early Stopping and KL Annealing

Early stopping is performed by keeping track of the best *𝓁*_total_ over the course of training. JAMIE is given a minimum number of epochs (default, 2, 500) over which time no early stopping can take place. Over this period, *κ* is gradually increased from 0 to 1. The concept originates from [27]. Once this period is over, the training continues until the algorithm does not produce a new best loss for a set number of epochs (default, 500) or until the maximum number of epochs is achieved (default, 10, 000). This process is further detailed in Algorithm 1.

### 4.8 JAMIE Algorithm

In Algorithm 1, datasets ***X*** and ***Y*** are provided pre-standardized along with an optional correspondence matrix ***F*** as described in Section 4. The user-given variables are *τ*_min_ (default, 2, 500) *τ*_max_ (default, 10, 000), which are the minimum and maximum number of epochs for training. *b*_*s*_ (default, 512) is the batch size. (*α, β, γ*) are the weights for the model with defaults (1000, 31.25, 31.25). *τ*_max lapses_ (default, 500) *𝓁*_min change_ (default, 1e-8) control the behavior of the early-stopping algorithm.

*ρ* is an experimental feature which controls how much the inferred correspondence matrix is utilized over the user-provided correspondence. The weighted average 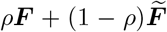 is used in Equation 3a and Equation 3b. By default, *ρ* = 1. This is not explored in-depth within the paper, but is a useful feature for low-correspondence applications.

### 4.9 Performance Evaluation

#### Label Transfer Accuracy (LTA)

To assess phenotype (e.g., cell type, cortical layer) separation, we use label transfer accuracy (LTA) [7,29]. LTA measures the phenotype-identification accuracy of a nearest neighbor classifier trained on the second modality and evaluated on the first. Higher values generally indicate better phenotype separation.

#### Fraction of Samples Closer to the True Match (FOSCTTM)

We use this metric to evaluate the average fraction of samples closer to the truly aligned sample (in both modalities) than its actual pairing. This provides an estimate of how closely the two modalities are aligned. Lower values generally indicate closer cross-modal alignment.

#### Pearson Correlation

We use Pearson correlation mainly to compare imputed and measured feature values, providing an estimate of how well the imputation method captures the variance of the measured feature values. For the purposes of imputation, higher values usually indicate better performance.

#### Area under the ROC curve (AUROC)

We use AUROC to evaluate imputed features for prediction of high or low measured expression values. To test this, we use AUROC on the median-binarized measured data, that is, each feature value above the feature median is treated as 1, and all others are treated as 0.

#### Jensen-Shannon Distance (JS distance)

It is also important that an imputation method preserves individual cell type distributions. We use JS distance, which is the square root of the Jensen-Shannon divergence. Jensen-Shannon divergence is the average KL divergence between two distributions and their average (a symmetrized version of KL divergence). KL divergence measures the difference between two probability distributions, and increases proportionally. As a baseline for imputation performance, a KNN model (*k* = 5) is used to generate new multimodal data based on the average of nearest neighbors in the training set.

##### Algorithm 1 JAMIE

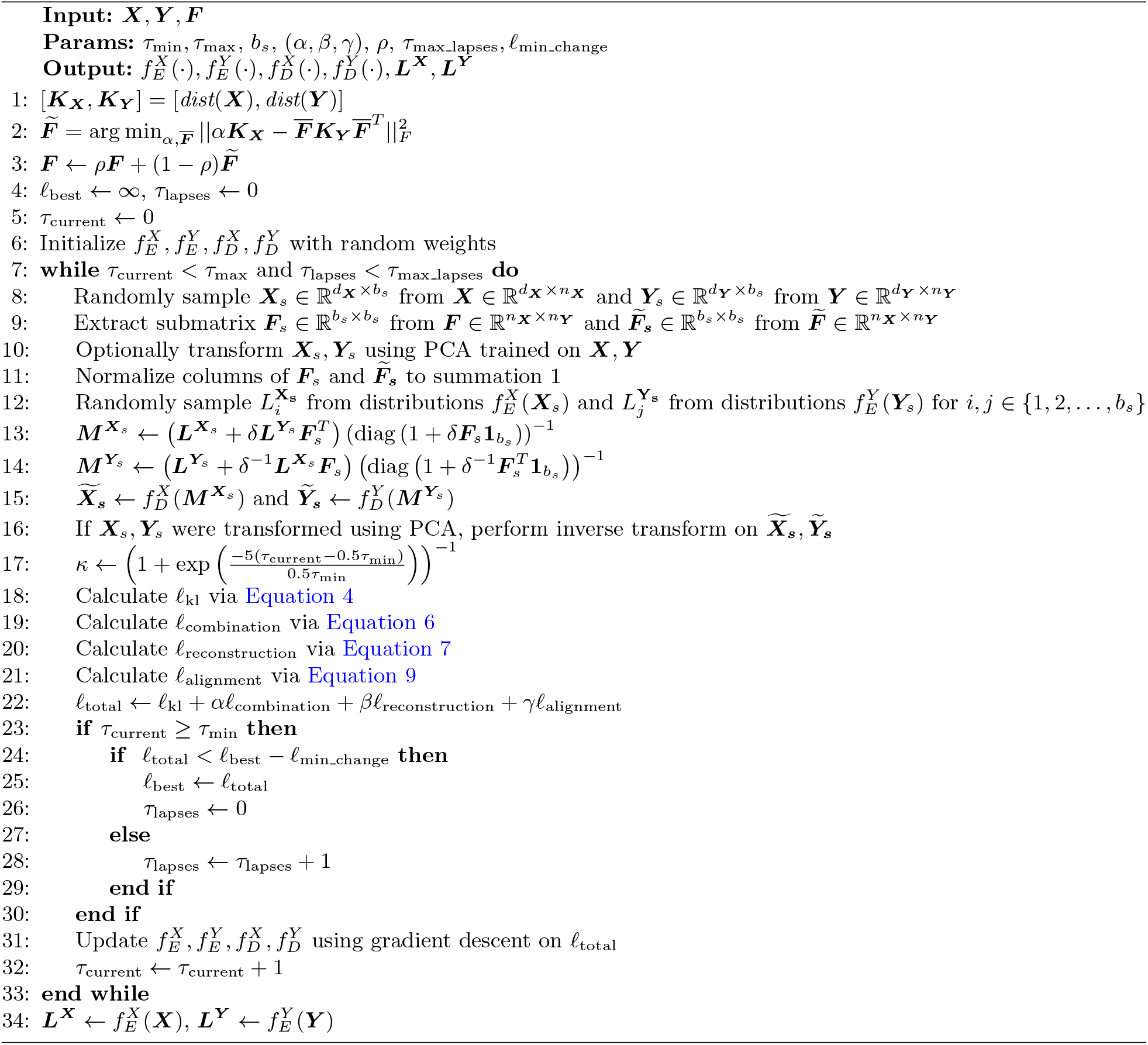

#### p-Values

When we compare two methods in terms of imputation performance, we utilize the two-tailed binomial test with the null hypothesis that both methods have an equal probability of imputing better on any given feature.

### 4.10 Hyperparameters and Validation

Principal Component Analysis (PCA) is used to lower the dimensionality of data with many features. The default value for this preprocessing is 512 features, and is automatically inverted when applying imputation. The hyperparameters used for all experiments in the paper are the default losses detailed in Section 4, the minimal and maximal iterations detailed in Subsection 4.7, and a batch size of 512.

When performing integration in Section 2, no data is withheld, hyperparameters are left at default, and dropout is set to *p* = 0 to facilitate maximal learning of single-modal nuances, regardless of dataset specificity. This is done because many alignment methods are unable to be reused, and perform integration only once before having to be recalculated. For imputation, hyperparameters are left unchanged but dropout is set at *p* = 0.6 to facilitate generalization. 20 percent of cells within each dataset are withheld at random. Any imputation performance calculations, such as those in Figure 3f, are performed on this withheld data.

### 4.11 Datasets and Preprocessing

We applied JAMIE to four main single-cell multimodal datasets: (1) simulated multimodal data generated by sampling from a Gaussian distribution on a branching manifold in MMD-MA (N=300, 3 cell types) [8]; (2) Patch-seq gene expression and electrophysiological features of single neuronal cells in the mouse visual cortex (N=3, 654, 6 cell types) [2] and mouse motor cortex (N=1, 208, 9 cell types) [30]; (3) 10x Multiome gene expression and chromatin accessibility data of 8, 981 cells in the human developing brain (21 postconceptional weeks, covering 7 major cell types in human cerebral cortex) [3]; (4) scRNA-seq gene expression and scATAC-seq chromatin accessibility data of 4,301 cells from the COLO-320DM colon adenocarcinoma cell line (which was also analyzed by BABEL [5]) [20]. The input datasets to JAMIE were standardized (mean = 0, standard deviation = 1) across features and large numbers of input features were reduced dimensionality by PCA to 512 PCs as needed for efficient computation.

### 4.12 Applications

#### Phenotype Prediction from Multimodal Data

Integration on multimodal datasets can improve classification, knowledge of phenotype, and understanding of complex biological mechanisms. Given two datasets ***X, Y*** and correspondence ***F***, we generate ***L***^***X***^, ***L***^***Y***^ from the trained encoders 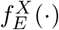, 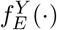 as in Subsection 4.1. Clustering can be performed or classification methods may be directly built atop the reusable latent space. Clustering on these latent spaces confers several advantages, the main of which being the incorporation of both modalities in the process of feature generation. We can then actively predict which samples correspond (if not already known) and perform related tasks such as cell type prediction. On partially-annotated datasets, cells in similar clusters should be of similar cell type. There is generally no need to use a complicated clustering or classification algorithm, as JAMIE should do most of the separation as a part of the latent space generation. The networks may be further analyzed to provide a clearer picture of the relationship between data features and phenotype. To visualize this cell type clustering, UMAP [31] is performed on high-dimensional data for visualization.

#### Cross-Modal Imputation

There are several methods to perform cross-modal imputation, but many methods are not able to prove that they have learned underlying biological mechanisms for the purpose of imputation. When utilized for cross-modal imputation, we can predict our missing data with a more rigorous foundation than that offered by a feed-forward network or linear regression. Given training data ***X, Y***, we can train the model. With new data 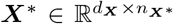, we can predict its correspondent latent embedding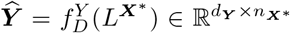 which has true values *Y* ^*\**^. Then, we have predicted correspondent cells using a latent space which likely confers understanding of the relationship between data features and phenotype.

This has motivation within the given losses. The alignment loss incentivizes the low-dimensional embeddings for each modality to be on the same latent space. Additionally, since we are predicting correspondent cells, ***F*** = 𝕀. Ideally then, ***L***^***X***^*\** = ***M***^***X***^*\** = ***M***^***Y***^ *\** = ***L***^***Y***^ *\**. So, if we assume perfect and total alignment, 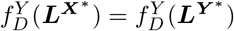. This prediction is shown in Figure 2b. Of course, this ideal scenario is not always the case. The efficacy of this cross-modal imputation method is explored within the results section of this paper, where 20 percent of the data is taken as validation, and the model is trained on the remainder.

#### Latent and Imputed Feature Interpretation

For interpretation of our trained model, we employ [17], henceforth referred to as SHAP. SHAP looks at individual predictions produced by a model and will assess importance of various input features through sample modulation. This can be used for a variety of interesting applications. If a target variable is easily separable by phenotype, SHAP can determine relevant features for further study. Additionally, if we perform imputation, SHAP can expose inter-modal linkages learned by the model. Given a model *f* and sample ***x*** *∈* ℝ^*n*^, SHAP values *ϕ*_0_, …, *ϕ*_*n−*1_ are learned such that 𝔼[*f* (*z*)] + Σ_*i=*{*j,…,m*}_*ϕ*_*i*_ = 𝔼[*f*(*z*) | *z*_*j*_ = *x*_*j*_,…,*z*_*m*_ = *x*_*m*_] for a background feature vector z ∈ R^*n*^. If {*j*, …, *m*} = {0, 1, …, *n −* 1}, then the sum of SHAP values and the background output will equal *f* (***x***), with each *ϕ*_*i*_ proportional to the impact of *x*_*i*_ on the model output.

Another useful technique involves choosing a key metric for classification (e.g., LTA) or imputation (e.g., Correspondence between imputed and measured features) and evaluating the metric with each feature sequentially removed (replaced with its background) from the model. Then, if the key metric becomes worse, it suggests that the removed feature is more important to the outcome of the model.

## 5 Data Availability

The MMD-MA simulation dataset can be downloaded from https://noble.gs.washington.edu/proj/mmd-ma/. Our simulation data may be downloaded from https://github.com/daifengwanglab/JAMIE. Processed Patch-seq gene expression and electrophysiological features for the mouse visual and motor cortices are available at https://github.com/daifengwanglab/scMNC. Raw Patch-seq datasets are available at [30,2]. Single-cell RNA-seq and ATAC-seq data on the human developing brain can be downloaded at https://github.com/GreenleafLab/brainchromatin/blob/main/links.txt under the heading *Multiome*. Single-cell RNA-seq and ATAC-seq of colon adenocarcinoma data can be found at https://github.com/wukevin/babel. Processed datasets for SNARE-seq adult mouse cortex data can be downloaded from https://scglue.readthedocs.io/en/latest/data.html.

## 6 Code Availability

All code was implemented in Python using PyTorch and the source code is publicly available at https://github.com/daifengwanglab/JAMIE [32]. Since Code Ocean provides an interactive platform for computational reproducibility [33], we have also provided an interactive version of our code for reproducing results and figures at [34].

## 7 Acknowledgments

This work was supported by National Institutes of Health grants R21NS128761, R21NS127432, R01AG067025 to D.W., P50HD105353 to Waisman Center, National Science Foundation Career Award 2144475 to D.W., and the start-up funding for D.W. from the Office of the Vice Chancellor for Research and Graduate Education at the University of Wisconsin–Madison. The funders had no role in study design, data collection and analysis, decision to publish, or manuscript preparation.

## 8 Author Contributions Statement

D.W. conceived and supervised the study. N.C.K. developed and implemented the methodology. X.H. and D.W. verified the methods. N.C.K. performed visualization and analysis. N.C.K, X.H., and D.W. edited and wrote the manuscript. All authors read and approved the final manuscript.

## 9 Competing Interests Statement

The authors declare no competing interests.

## Notes

### Competing Interest Statement

The authors have declared no competing interest.

### Summary of Updates

We fixed math typos and clarified some text.

