## Supplemental Materials for "Joint Variational Autoencoders for Multimodal Imputation and Embedding"

### 1 Method Comparison

JAMIE has a wide selection of features compared to existing methods, and is flexible in its implementation. A brief overview of features and applications can be seen in Table 1. More details can be found in Section 4 and Subsection 4.12.

**Supplementary Table 1.** Method comparison between various integration and imputation methods. JAMIE unifies features from several different integration and imputation methods in a single architecture. NLMA: nonlinear manifold alignment [1], UnionCom [2], CCA: canonical correlation analysis [1, 3], BABEL [4].

| | JAMIE | N/LMA | UnionCom | MMD-MA | CCA | BABEL |
|---|---|---|---|---|---|---|
| Cell Type Separation | ✓ | ✓ | ✓ | ✓ | ✓ | |
| Generalizable Model | ✓ | | | | ✓ | ✓ |
| Latent Space | ✓ | ✓ | ✓ | ✓ | ✓ | ✓ |
| Missing Modality Imputation | ✓ | | | | | ✓ |
| Model Type | | | | | | |
| Autoencoder | ✓ | | | | | ✓ |
| Eigenanalysis | | ✓ | | | ✓ | |
| Manifold Alignment | | ✓ | | ✓ | | |
| Projection | | | ✓ | | | |
| Non-Omics Data Compatibility | ✓ | | | ✓ | | |
| Partial Correspondence | ✓ | ✓ | | | ✓ | |

### 2 Datasets

MMD-MA provides simulation data which produces a branching pattern when mapped into 2d space [5]. The dataset has data matrices $\mathbb{R}^{300\times 1,000}, \mathbb{R}^{300\times 2,000}$ over 3 cell types.

We generated simulation data using scMultiSim for RNA and ATAC sequencing using the default package setting `GRN_params_1139` [6]. The dataset contains 10 cell types and measures $\mathbb{R}^{500\times 1,250}, \mathbb{R}^{500\times 3,750}$ for RNA and ATAC, respectively.

We use data of the mouse visual cortex measured with patch-seq, which profiles gene expression and electrophysiological data of neuronal cells [7]. The data contains $3,654$ cells of 6 distinct types. Compatibility with non-omics data is important to the mission of JAMIE, which is why this dataset is invaluable for testing. The data matrices measure $\mathbb{R}^{3,654\times 1,302}$ for gene expression and $\mathbb{R}^{3,654\times 39}$ for electrophysiological features.

Mouse motor cortex data uses patch-seq to measure gene expression and electrophysiological data of neuronal cells [8]. The data matrices measure $\mathbb{R}^{1,208\times 1,286}$ for gene expression and $\mathbb{R}^{1,208\times 29}$ for electrophysiological features. Each cell is classified as one of 9 distinct cell types, characterized by a marker gene.

Developing brain data contains scRNA-seq and scATAC-seq data [9]. The data contains $8,981$ cells of 7 types in the human developing brain (21 postconceptional weeks), categorized by the top cis-regulatory element (CRE) linked genes. The data matrices measure $\mathbb{R}^{8,981\times 34,104}$ for RNA and $\mathbb{R}^{8,981\times 19,836}$ for ATAC, and demonstrates the capabilities of JAMIE on raw data.

We also use scRNA-seq and scATAC-seq data from the COLO-320DM cell line sequencing colon adenocarcinomas from a single subject [4, 10]. The data measures $\mathbb{R}^{4,301\times 34,861}$ for RNA and $\mathbb{R}^{4,301\times 85,596}$ for ATAC. This dataset was utilized in [4], and serves as a benchmark for imputation performance.

Lastly, we use scRNA-seq and scATAC-seq data from SNARE-seq on the adult mouse cerebral cortex [11]. The data matrices measure $\mathbb{R}^{9,190\times 28,930}$ for RNA and $\mathbb{R}^{9,190\times 241,757}$ for ATAC. The dataset was utilized for testing in [12].

### 3 Hyperparameters for Supporting Methods

All comparison methods use default hyperparameters for each dataset. In particular, NLMA and LMA [1] use $\mu = 0.9, \epsilon = 10^{-8}$, UnionCom [2] runs $2,000$ iterations of its prime-dual method and 100 for computing the deep neural network with parameters $\mathrm{lr} = 0.001$ and batch size 100. MMD-MA [5] uses $\mathrm{lr} = 5\times 10^{-5}$ and default tradeoff parameters over $10,000$ iterations. The kernel matrices are calculated via multiplication of the data matrix with its transpose. We also used the default parameters provided by scGLUE [12]: dropout 0.2 with 2 hidden layers of 256 nodes each for the encoder and decoders. We also follow the documentation for constructing the prior regulatory graph from the data preprocessed by scGLUE [12]. BABEL [4] uses batch size 512 and learning rate 0.01.

**Supplementary Table 2.** Runtime for each dataset at *Preparation* and *Training* phases. *Preparation* is the calculation of $\widetilde{\boldsymbol{F}}$ using the method described in Subsection 4.6. *Training* is the primary training phase of JAMIE where the dual variational autoencoders are generated. **Bolding** in the memory column indicates that the memory usage was dominated by the intra-dataset distance calculation. In the runtime column, this indicates which phase took the most time.

| Dataset | | Feature Numbers | | Usage | | Runtime (s) | |
|---|---|---|---|---|---|---|---|
| Citation | Cells | Modal 1 | Modal 2 | Epochs | Memory (Mb) | Preparation | Training |
| Liu et al. [5] | 300 | 2,000 | 1,000 | 3,800 | 17 | 4 | **107** |
| Li et al. [6] | 500 | 1,250 | 3,750 | 6,600 | 59 | 12 | **470** |
| Scala et al. [8] | 1,208 | 1,286 | 29 | 3,800 | 42 | 173 | **353** |
| Gouwens et al. [7] | 3,654 | 1,302 | 39 | 3,400 | **319** | **4,508** | 1,122 |
| Wu et al. [4] | 4,301 | 34,861 | 85,596 | 3,300 | **5,160** | **7,377** | 2,188 |
| Trevino et al. [9] | 8,981 | 34,104 | 19,836 | 3,100 | 4,846 | **44,687** | 4,686 |
| Chen, Lake, and Zhang [11] | 9,190 | 28,930 | 241,757 | 3,400 | **30,442** | **47,500** | 5,057 |

### 4 Supplementary Figures

Visualizations were created with the help of [13] for automatically placing labels and [14] for facilitating axis discontinuities. For various runs of JAMIE, not all prior correspondence information is provided. In figures, we refer to these based on the percentage of correspondent multimodal pairings provided to JAMIE. e.g., 50% provided correspondence becomes (.5) and 75% provided correspondence becomes (.75). For an example, see Supplementary Figure 4.

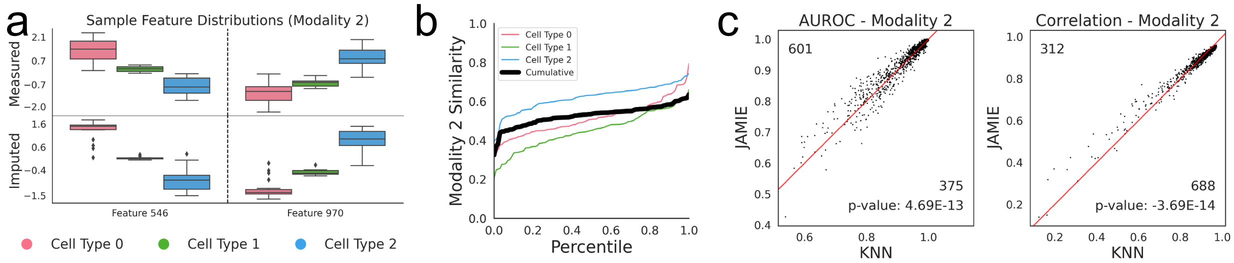

**Supplementary Figure 1.** JAMIE imputation of simulated Gaussian data on a branching manifold [5]. (a) Select features showing measured (top) and imputed (bottom) features by cell type ($n = 300$). Boxes span from the upper to lower quartiles and a line indicates the median. Whiskers extend to extremes up to 1.5 times the interquartile range from the upper and lower quartiles. Any outliers beyond this range are plotted individually. (b) Similarity (1 − JS distance) between measured and imputed value distributions of the second modality. The black line indicates the average similarity across cell types. (c) Imputation performance of JAMIE versus a baseline KNN for AUROC and correlation. We utilize a two-tailed binomial test to generate p-values.

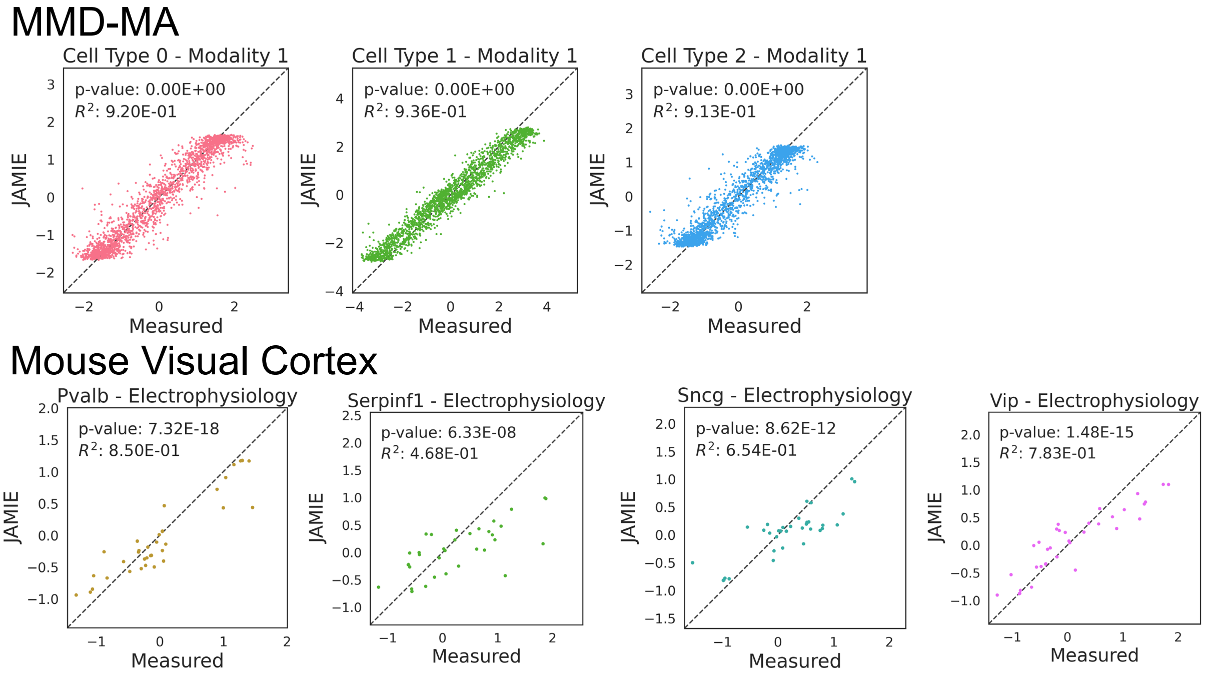

**Supplementary Figure 2.** Measured (x-axis) vs imputed (y-axis) data for select cell types, including $R^2$ and correlation $p$-values. We utilize Pearson correlation to generate p-values. Top: Liu et al. [5]. Bottom: Gouwens et al. [7].

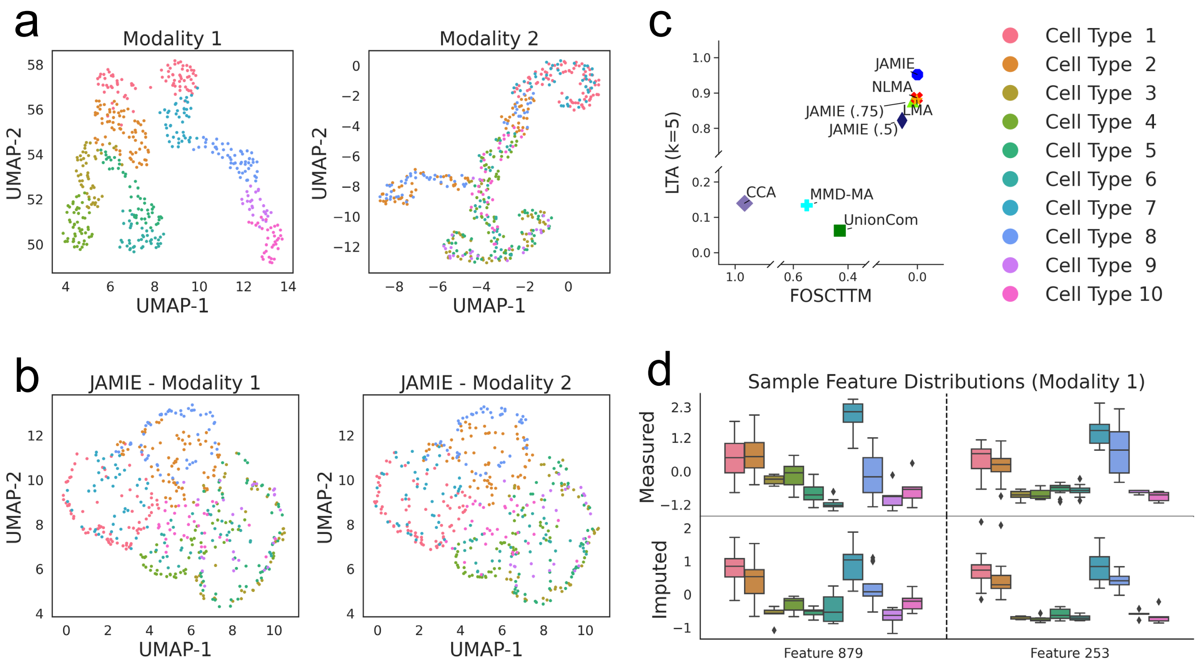

**Supplementary Figure 3.** JAMIE integration performance on simulated RNA and ATAC-seq data from scMultiSim [6] as detailed in Supplementary Section 2. (a) UMAP of single cells by gene expression data (left) and chromatin accessibility data (right), colored by cell types. (b) JAMIE latent spaces of (a). (c) Fraction of samples closer to the true mean (x-axis) and label transfer accuracy (y-axis) of JAMIE and state-of-the-art methods for cell type separation using all correspondence information available. CCA: canonical correlation analysis [3, 1], LMA: linear manifold alignment [1], MMD-MA [5], NLMA: nonlinear manifold alignment [1], UnionCom [2]. (d) Measured (top) and imputed (bottom) values of two chromatin accessibility features across cell types ($n = 1,250$). Boxes span from the upper to lower quartiles and a line indicates the median. Whiskers extend to extremes up to 1.5 times the interquartile range from the upper and lower quartiles. Any outliers beyond this range are plotted individually.

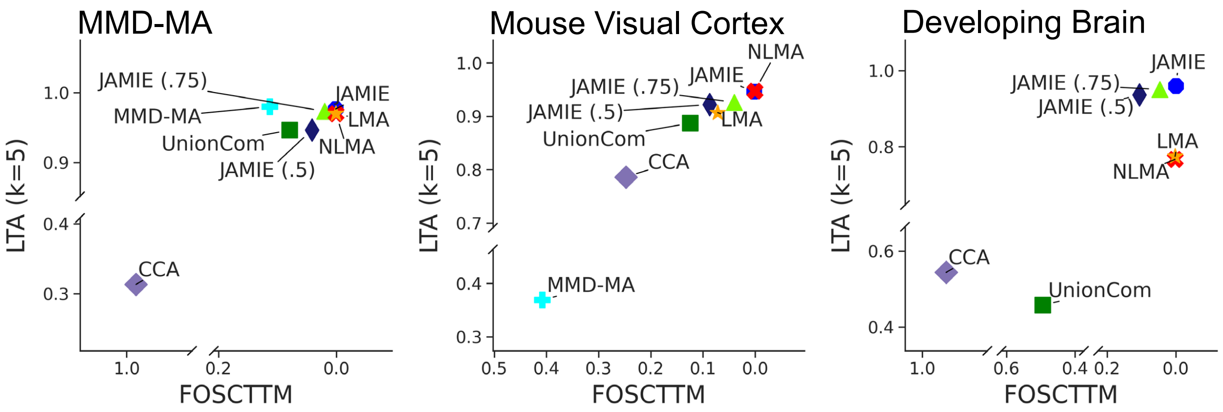

**Supplementary Figure 4.** Extended panel (c) for Figure 3, Figure 4, and Figure 5, respectively, including JAMIE run with only 75% (.75) and 50% (.5) correspondence information provided. Fraction of samples closer to the true mean (x-axis) and label transfer accuracy (y-axis) of JAMIE with full and partial correspondence provided and state-of-the-art methods for cell type separation using all correspondence information available. CCA: canonical correlation analysis [3, 1], LMA: linear manifold alignment [1], MMD-MA [5], NLMA: nonlinear manifold alignment [1], UnionCom [2].

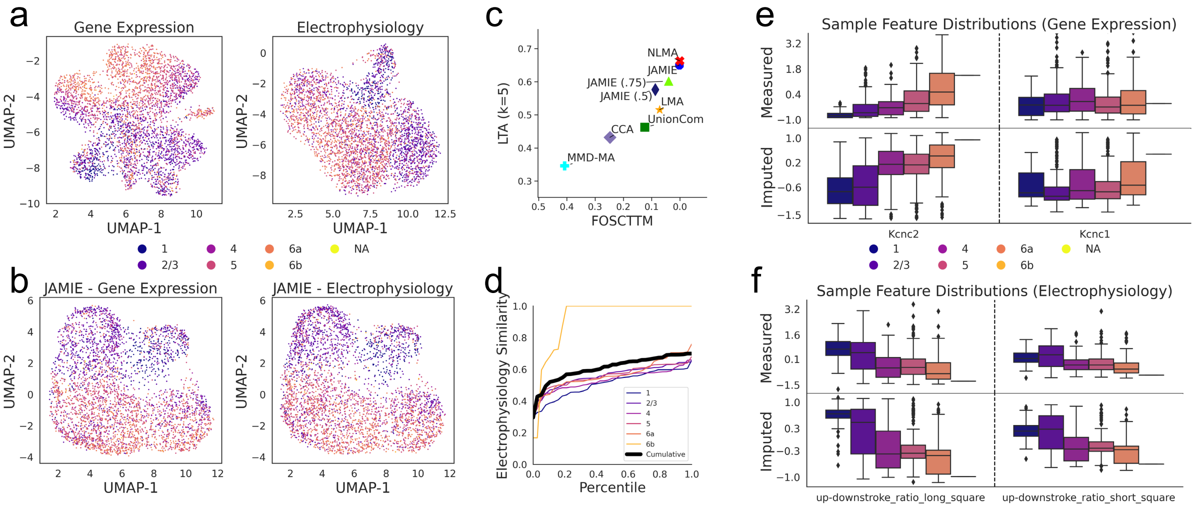

**Supplementary Figure 5.** Gene expression and electrophysiological features in the mouse visual cortex [7]. (a) UMAP of single cells by gene expression data (left) and electrophysiological data (right), colored by cortical layer. (b) JAMIE latent spaces of (a). (c) Fraction of samples closer to the true mean (x-axis) and label transfer accuracy (y-axis) of JAMIE with full and partial correspondence provided and state-of-the-art methods for cell type separation using all correspondence information available. CCA: canonical correlation analysis [3, 1], LMA: linear manifold alignment [1], MMD-MA [5], NLMA: nonlinear manifold alignment [1], UnionCom [2]. (d) Cumulative distributions of similarity ($1 - \text{JS distance}$) between measured and imputed electrophysiological features by gene expression. The black line indicates the average similarity across cell types. (e) Measured (top) and imputed (bottom) values of two gene expression features across cell types ($n = 3,624$). Boxes span from the upper to lower quartiles and a line indicates the median. Whiskers extend to extremes up to 1.5 times the interquartile range from the upper and lower quartiles. Any outliers beyond this range are plotted individually. (f) Same as (e) for electrophysiological features.

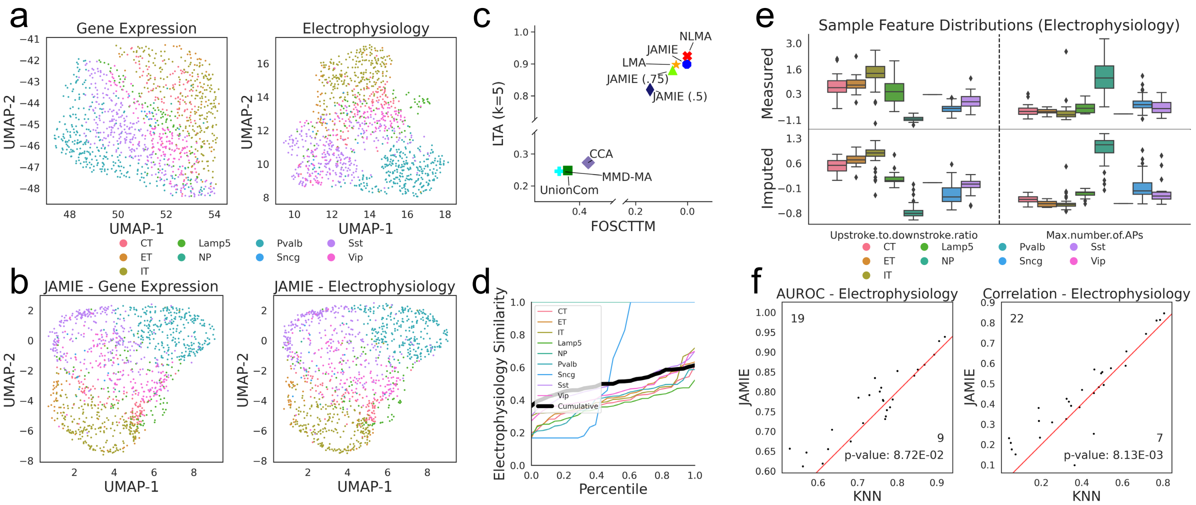

**Supplementary Figure 6.** Gene expression and electrophysiological features in the mouse motor cortex [8]. (a) UMAP of single cells by gene expression data (left) and electrophysiological data (right), colored by cell types (inhibitory neuronal types). (b) JAMIE latent spaces of (a). (c) Fraction of samples closer to the true mean (x-axis) and label transfer accuracy (y-axis) of JAMIE with full and partial correspondence provided and state-of-the-art methods for cell type separation using all correspondence information available. CCA: canonical correlation analysis [3, 1], LMA: linear manifold alignment [1], MMD-MA [5], NLMA: nonlinear manifold alignment [1], UnionCom [2] (d) Cumulative distributions of similarity ($1-$ JS distance) between measured and imputed electrophysiological features by gene expression. The black line indicates the average similarity across cell types. (e) Measured (top) and imputed (bottom) values of two electrophysiological features across cell types ($n = 1,208$). Boxes span from the upper to lower quartiles and a line indicates the median. Whiskers extend to extremes up to 1.5 times the interquartile range from the upper and lower quartiles. Any outliers beyond this range are plotted individually. (f) Performance for imputing electrophysiological features of JAMIE versus a baseline KNN by AUROC and correlation. We utilize a two-tailed binomial test to generate p-values.

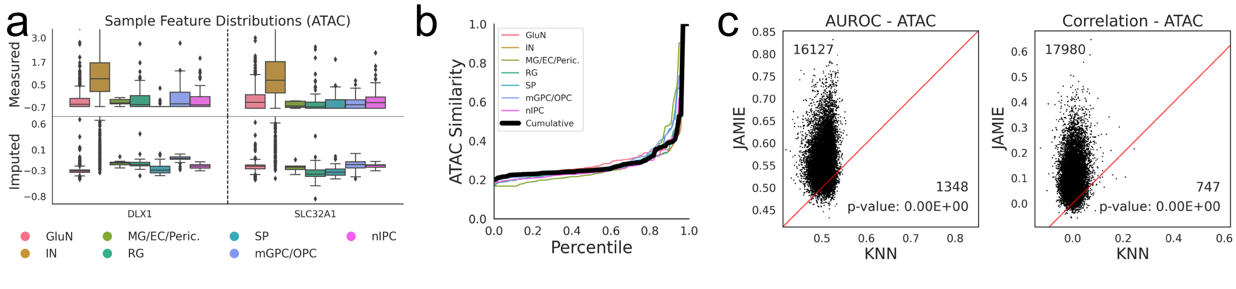

**Supplementary Figure 7.** Chromatin accessibility imputed from gene expression in the developing brain [9]. (a) Select peaks showing measured (top) and imputed (bottom) chromatin accessibility by cell type ($n = 8,981$). Boxes span from the upper to lower quartiles and a line indicates the median. Whiskers extend to extremes up to 1.5 times the interquartile range from the upper and lower quartiles. Any outliers beyond this range are plotted individually. (b) Similarity ($1-$ JS distance) between measured and imputed chromatin accessibility distributions. The black line indicates the average similarity across cell types. (c) Imputation performance of JAMIE versus a baseline KNN for AUROC and correlation. We utilize a two-tailed binomial test to generate p-values.

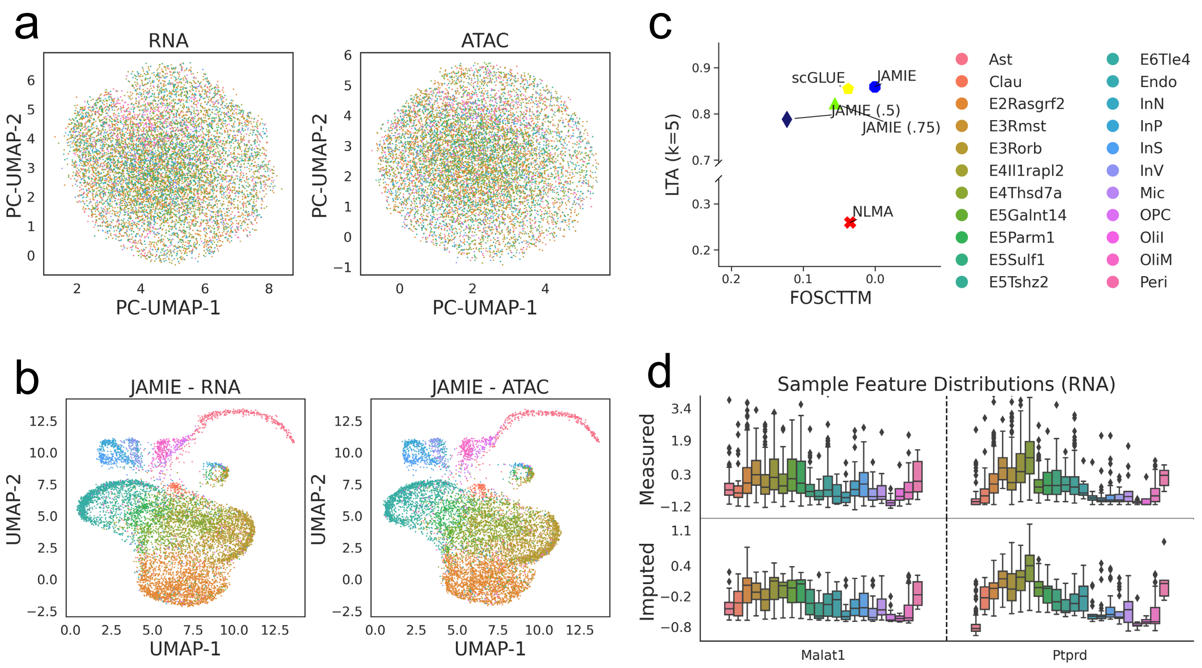

**Supplementary Figure 8.** JAMIE application and benchmarking using SNARE-seq data (gene expression and chromatin accessibility) on the adult mouse cerebral cortex [11]. (a) UMAP of 4,096 principal components of single cells by gene expression data (left) and chromatin accessibility data (right), colored by cell types. (b) JAMIE latent spaces of (a). (c) Fraction of samples closer to the true mean (x-axis) and label transfer accuracy (y-axis) of JAMIE and state-of-the-art methods for cell type separation using all correspondence information available. scGLUE: graph-linked unified embedding [12], NLMA: nonlinear manifold alignment [1]. CCA [3, 1] collapsed during runtime and is not included for clarity. (d) Measured (top) and imputed (bottom) values of two chromatin accessibility features across cell types ($n = 9,190$). Boxes span from the upper to lower quartiles and a line indicates the median. Whiskers extend to extremes up to 1.5 times the interquartile range from the upper and lower quartiles. Any outliers beyond this range are plotted individually.

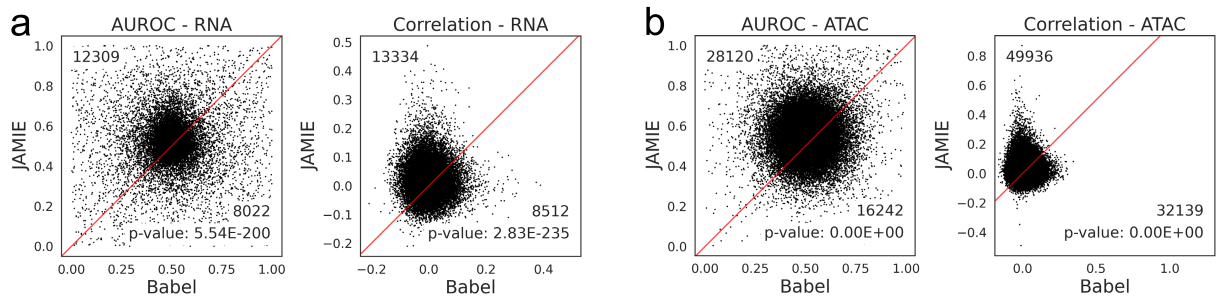

**Supplementary Figure 9.** JAMIE and BABEL [4] compared by imputation performance on colon adenocarcinomas for (a) ATAC to RNA and (b) RNA to ATAC imputation. AUROC and correlation by feature are shown for each comparison. Points are only plotted for defined values. We utilize a two-tailed binomial test to generate p-values.

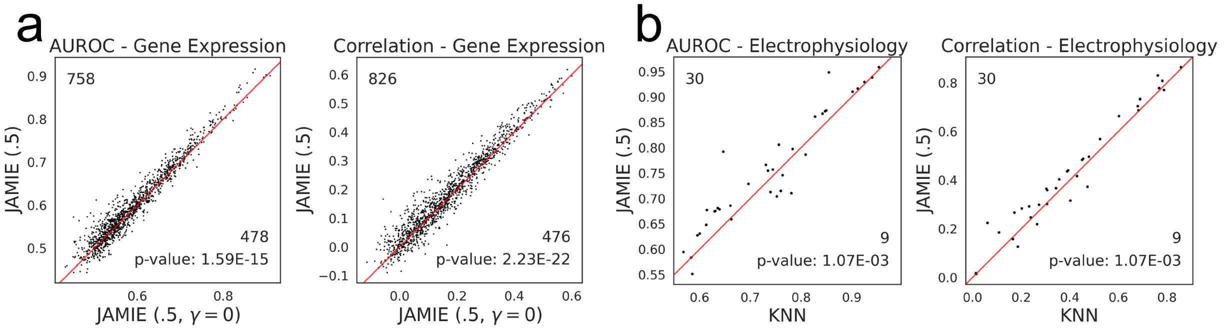

**Supplementary Figure 10.** Imputation performance using 50% of user-provided correspondence on gene expression and electrophysiological features in the mouse visual cortex [7]. (a) Performance for imputing gene expression features of JAMIE using 50% of user-provided correspondence versus JAMIE without alignment loss by AUROC and correlation. (b) Performance for imputing electrophysiological features of JAMIE using 50% of user-provided correspondence versus a baseline KNN using 100% of user-provided correspondence by AUROC and correlation. For both subfigures, we utilize a two-tailed binomial test to generate p-values.

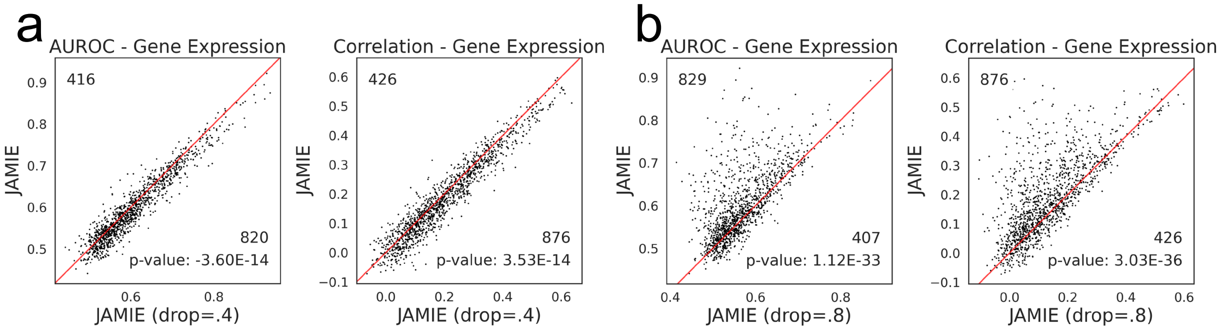

**Supplementary Figure 11.** Imputation performance using differing dropout values on gene expression and electrophysiological features in the mouse visual cortex [7]. (a) Performance for imputing gene expression features of JAMIE using dropout $p = 0.4$ versus $p = 0.6$ (default) by AUROC and correlation. $p = 0.4$ achieved a mean AUROC and correlation of 0.607 and 0.199 for imputing gene expression, with 0.734 and 0.475 for imputing electrophysiological features. $p = 0.6$ achieved a mean AUROC and correlation of 0.598 and 0.183 for imputing gene expression, with 0.761 and 0.470 for imputing electrophysiological features. (b) Performance for imputing gene expression features of JAMIE using dropout $p = 0.8$ versus $p = 0.6$ (default) by AUROC and correlation. $p = 0.8$ achieved a mean AUROC and correlation of 0.566 and 0.131 for imputing gene expression, with 0.713 and 0.425 for imputing electrophysiological features. For both subfigures, we utilize a two-tailed binomial test to generate p-values.

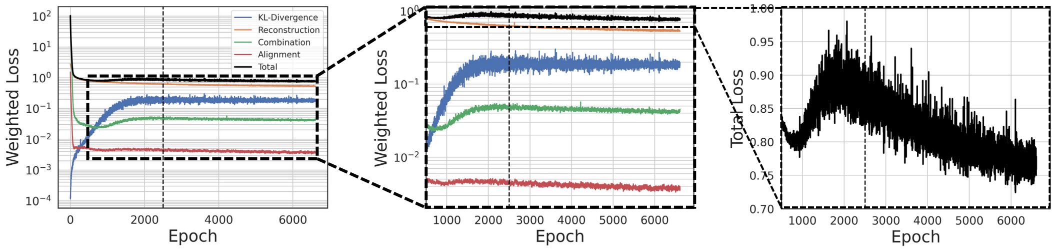

**Supplementary Figure 12.** Weighted losses over the course of training on simulated RNA and ATAC-seq data from scMultiSim [6] as detailed in Supplementary Section 2. The vertical black dotted line indicates the end of KL-annealing.

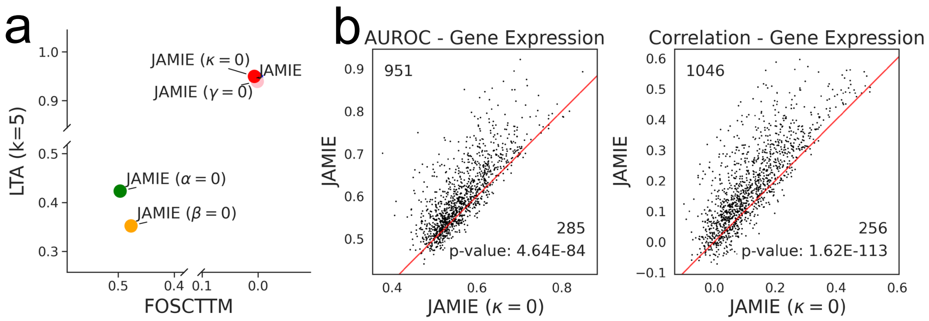

**Supplementary Figure 13.** Training loss comparison on gene expression and electrophysiological features in the mouse visual cortex [7]. (a) Fraction of samples closer to the true mean (x-axis) and label transfer accuracy (y-axis) of JAMIE with each loss individually removed. (b) Performance for imputing gene expression of JAMIE with and without $\ell_{\mathrm{kl}}$ (Subsection 4.3) by AUROC and correlation. We utilize a two-tailed binomial test to generate p-values.

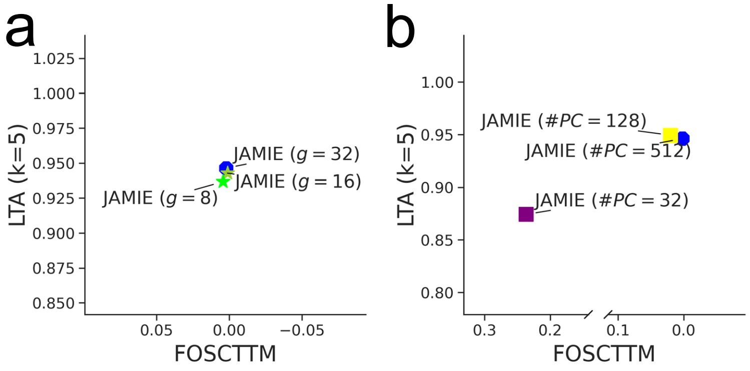

**Supplementary Figure 14.** Resultant latent space quality for varying latent dimensions and number of principal components ($\#PC$) on gene expression and electrophysiological features in the mouse visual cortex [7]. (a) JAMIE for different latent space dimensions $g$. (b) JAMIE for different numbers of principal components.

[7] Nathan W Gouwens, Staci A Sorensen, Fahimeh Baftizadeh, Agata Budzillo, Brian R Lee, Tim Jarsky, Lauren Alfiler, Katherine Baker, Eliza Barkan, Kyla Berry, Darren Bertagnolli, Kris Bickley, Jasmine Bomben, Thomas Braun, Krissy Brouner, Tamara Casper, Kirsten Crichton, Tanya L Daigle, Rachel Dalley, Rebecca A de Frates, Nick Dee, Tsega Desta, Samuel Dingman Lee, Nadezhda Dotson, Tom Egdorf, Lauren Ellingwood, Rachel Enstrom, Luke Esposito, Colin Farrell, David Feng, Olivia Fong, Rohan Gala, Clare Gamlin, Amanda Gary, Alexandra Glandon, Jeff Goldy, Melissa Gorham, Lucas Graybuck, Hong Gu, Kristen Hadley, Michael J Hawrylycz, Alex M Henry, Dijon Hill, Madie Hupp, Sara Kebede, Tae Kyung Kim, Lisa Kim, Matthew Kroll, Changkyu Lee, Katherine E Link, Matthew Mallory, Rusty Mann, Michelle Maxwell, Medea McGraw, Delissa McMillen, Alice Mukora, Lindsay Ng, Lydia Ng, Kiet Ngo, Philip R Nicovich, Aaron Oldre, Daniel Park, Hanchuan Peng, Osnat Penn, Thanh Pham, Alice Pom, Zoran Popović, Lydia Potekhina, Ramkumar Rajanbabu, Shea Ransford, David Reid, Christine Rimorin, Miranda Robertson, Kara Ronellenfitch, Augustin Ruiz, David Sandman, Kimberly Smith, Josef Sulc, Susan M Sunkin, Aaron Szafer, Michael Tieu, Amy Torkelson, Jessica Trinh, Herman Tung, Wayne Wakeman, Katelyn Ward, Grace Williams, Zhi Zhou, Jonathan T Ting, Anton Arkhipov, Uygar Sümbül, Ed S Lein, Christof Koch, Zizhen Yao, Bosiljka Tasic, Jim Berg, Gabe J Murphy, and Hongkui Zeng. “Integrated Morphoelectric and Transcriptomic Classification of Cortical GABAergic Cells”. en. In: *Cell* 183.4 (Nov. 2020), 935–953.e19.
